# Skeletal muscle hypertrophy rewires glucose metabolism: an experimental investigation and systematic review

**DOI:** 10.1101/2022.12.08.519580

**Authors:** Philipp Baumert, Sakari Mäntyselkä, Martin Schönfelder, Marie Heiber, Mika Jos Jacobs, Anandini Swaminathan, Petras Minderis, Mantas Dirmontas, Karin Kleigrewe, Chen Meng, Michael Gigl, Ildus I. Ahmetov, Tomas Venckunas, Hans Degens, Aivaras Ratkevicius, Juha J. Hulmi, Henning Wackerhage

## Abstract

**Background:** Proliferating cancer cells shift their metabolism toward glycolysis even in the presence of oxygen to especially generate glycolytic intermediates as substrates for anabolic reactions. We hypothesize that a similar metabolic remodeling occurs during skeletal muscle hypertrophy.

**Methods:** We used mass spectrometry in hypertrophying C2C12 myotubes *in vitro* and plantaris mice muscle *in vivo* and assessed metabolomic changes and the incorporation of [U-^13^C_6_]glucose tracer. We performed enzyme inhibition of the key serine synthesis pathway enzyme phosphoglycerate dehydrogenase (Phgdh) for further mechanistic analysis and conducted a systematic review to align any changes in metabolomics during muscle growth with published findings. Finally, UK Biobank was used to link the findings to population level.

**Results:** The metabolomics analysis in myotubes revealed IGF-1 induced altered metabolite concentrations in anabolic pathways such as in the pentose phosphate (ribose-5-phosphate/ribulose-5-phosphate: +40%; p=0.01) and serine synthesis pathway (serine: - 36.8%; p=0.009). Like the hypertrophy-stimulation with IGF-1 in myotubes *in vitro*, the concentration of the dipeptide L-carnosine was decreased by 26.6% (p=0.001) during skeletal muscle growth *in vivo.* However, phosphorylated sugar (glucose-6-phosphate, fructose-6-phosphate or glucose-1-phosphate) decreased by 32.2% (p=0.004) in the overloaded muscle *in vivo*, while increased in the IGF-1 stimulated myotubes *in vitro*. The systematic review revealed that 10 metabolites linked to muscle hypertrophy were directly associated with glycolysis and its interconnected anabolic pathways. We demonstrated that labelled carbon from [U-^13^C_6_]glucose is increasingly incorporated by ∼13% (p=0.001) into the non-essential amino acids in hypertrophying myotubes, which is accompanied by an increased depletion of media serine (p=0.006). The inhibition of Phgdh suppressed muscle protein synthesis in growing myotubes by 58.1% (p<0.001) highlighting the importance of the serine synthesis pathway for maintaining muscle size. Utilizing data from the UK Biobank (n=450,243), we then discerned genetic variations linked to the serine synthesis pathway (*PHGDH* and *PSPH*) and to its downstream enzyme (*SHMT1*), revealing their association with appendicular lean mass in humans (p<5.0e-8).

**Conclusion:** Understanding the mechanisms that regulates skeletal muscle mass will help in developing effective treatments against muscle weakness. Our results provide evidence for metabolic rewiring of glycolytic intermediates into anabolic pathways during muscle growth, such as in the serine synthesis.

## INTRODUCTION

Muscle hypertrophy and increased strength are typical phenotypic adaptations to resistance training or chronic muscle overload in mice [1]. Resistance training is an effective low-cost intervention for the prevention and treatment of muscle loss in healthy and diseased individuals [2]. Resistance training not only increases muscle size and strength but also improves glycemic control, increases glucose uptake as well as insulin sensitivity, suggesting metabolic adaptations in healthy as well as diabetic persons. While mechanistic target of rapamycin complex 1 (mTORC1) has been identified as a key mediator of the increased protein synthesis after resistance exercise and during chronic overload [3], there are still many gaps in our understanding of how changes in the muscle metabolism possibly contributes to muscle hypertrophy [1] or atrophy [4].

Research, especially on cancer, shows that proliferating cells reprogram their metabolism and synthesize lactate in the presence of oxygen, which has been termed the Warburg effect. One key function of the Warburg effect is to synthesize glycolytic metabolites as substrates for anabolic reactions that build biomass [5]. About 20% of the carbon dry mass of cancer cells is delivered from glucose carbon [6], which is accomplished by directing some glycolytic intermediates toward anabolic pathways such as the serine synthesis and pentose phosphate pathway. These two pathways generate critical precursors (e.g., non-essential amino acids and nucleotides) for the synthesis of macromolecules that are required for cancer cell proliferation [7, 8]. Conversely, knockdown of the key enzyme phosphoglycerate dehydrogenase (Phgdh), which diverts the glucose-derived 3-phosphoglycerate away from glycolysis into the *de novo* serine synthesis pathway, reduces cancer cell proliferation [9] indicating the importance of this pathway for cellular growth and proliferation. We recently hypothesized that a Warburg-like metabolic reprogramming also occurs in mature, hypertrophying muscle fibers [10]. Consistent with this, several investigations indicate increased expression of genes that encode enzymes of anabolic pathways [11, 12], and a rewiring of glucose metabolism in hypertrophying muscles [13–16].

In this study, by using a mass spectrometry-based metabolomics approach in hypertrophy-stimulated muscles both in C2C12 myotubes *in vitro* and in plantaris mice muscle *in vivo*, we identified metabolite level changes linked to the pentose phosphate, and the serine synthesis pathways in hypertrophy-stimulated muscles. We additionally analyzed pre-existing metabolomics datasets in mice and utilized the UK Biobank to identify genetic variations and their association with human appendicular lean mass further indicating a role of metabolic pathways parallel to glycolysis in muscle growth. Stable isotope [U-^13^C_6_]glucose tracer revealed that carbon of glucose is increasingly incorporated into non-essential amino acids during muscle growth. Finally, we inhibited the key enzyme of the serine synthesis pathway in differentiated muscle cells revealing an important role of this pathway in protein synthesis. Our results further demonstrate that a hypertrophying skeletal muscle reprograms its metabolism to support biomass production.

## MATERIAL AND METHODS

### Animals and *in vivo* study design

All procedures involving mice were approved by the Lithuanian State Food and Veterinary Service (Animal ethics number: 2018-06-14 Nr. G2-90). Male C57BL/6J mice were used in all experiments. Mice were housed in standard cages, one to three mice per cage at a temperature of 22–24°C and 40–60% humidity. Animals were fed standard chow diet and received tap water ad libitum.

Six mice were subjected to muscle overload of the plantaris muscle for 14 days starting at the age of 32 weeks. After cessation of nociceptive responses, an incision was made in the popliteal area of the left hind limb to expose the tibial nerve with blunt dissection. The branches of the tibial nerve that innervate the soleus and gastrocnemius muscles were cut and small segments removed to prevent reinnervation. This strategy imposes an overload and subsequent compensatory hypertrophy of the plantaris muscle. The contralateral plantaris muscle of these mice served as internal control in analysis of muscle mass and metabolomics measurements. After 14 days, mice were fasted overnight, and they were sacrificed by the exposure to CO_2_. Immediately after sacrifice, the plantaris muscle was excised, blotted dry, weighed and frozen in isopentane (pre-cooled with liquid nitrogen). The muscle was then stored at −80 °C until further metabolomics analysis. For a detailed description, see supporting information.

### Cell culture and *in vitro* study design

Mouse C2C12 myoblasts cells were cultured in Dulbecco’s modified Eagle’s medium (DMEM) supplemented with 10% fetal bovine serum. To induce differentiation, the myoblasts were cultured until confluence and then the medium was switched to DMEM with 0.2% FBS. After 72 h of differentiation, C2C12 Myotubes were serum starved (DMEM only) for 4 h and then treated with DMEM including 0.2% dialyzed FBS and vehicle control (10 mM HCL, 0.1% bovine serum albumin; BSA), insulin-like growth factor 1 (IGF-1; 100 ng/ml; recombinant Human LONG R^3^ IGF-1, Sigma-Aldrich, Cat# I1271, MO, USA), rapamycin (10 ng/ml; Calbiochem, Cat#553210, Watford, Hertfordshire, UK) and IGF-1 + rapamycin, for 48 h. For Phgdh inhibition experiments, we treated C2C12 cells for 48 h with chemical PHGDH inhibitor NCT-503 (50 µM; MCE, HY-101966; MedChemExpress) five days after the initiation of differentiation [17]. To measure carbon incorporation in IGF-1 stimulated C2C12 myotubes, 4.5 g/L glucose consisting of ∼20% [U-^13^C_6_]glucose and of 80% non-labelled glucose was added into glucose-free DMEM supplemented with 2 % horse serum (Sigma-Aldrich, H1138, NY, USA). Cells were incubated for 24 h with isotope-enriched media on day 3 post differentiation after which the media was collected. Finally, cells were harvested in 1 M perchloric acid (PCA) for stable isotope analysis. Please see the Supporting Information for detailed experimental procedures.

### RNA isolation, reverse transcription, and quantitative real-time PCR

C2C12 myotubes were washed twice with PBS at room temperature and then lysed in lysis T buffer (Peqlab Biotechnology GmbH, reference number 12-6634-01). Total RNA was then extracted with the peqGOLD Total RNA Kit C-Line (Peqlab Biotechnology GmbH, reference number: 12-6634-01) according to the manufacturer’s instructions. For mRNA analysis, we synthesised cDNA using 2 µg of total RNA and cDNA was amplified with a PerfeCTa qPCR fluorescent SYBR Green SuperMix (Quantabio) using real-time quantitative PCR (Rotor-Gene RG 6000— QIAGEN). The primer sequences used for the gene expression are shown in Table S1. Please see the Supporting Information for detailed experimental procedures.

### Protein determination and western blot analysis

Cells were lysed on ice in 100 µl of radioimmunoprecipitation assay buffer (RIPA) including 0.1% SDS, 0.5 M sodium orthovanadate, 0.5% sodium deoxycholate, 50 mM NaF, 1 mM EDTA,150 mM NaCl, 1% Triton-X 100 and with 1X protease and phosphatase inhibitor cocktail (Peqlab Biotechnology GmbH, Germany). For the Phgdh inhibition experiment, protein synthesis in C2C12 myotubes was measured using SUnSET protein synthesis measurement. Specific protein quantification was performed by western blot analysis. Protein samples containing 40 µg protein were solubilized in Laemmli sample buffer containing 10% β-mercaptoethanol and heated for 5 minutes at 95°C. Samples were then separated by electrophoresis in a 10% SDS-PAGE gel (Bio-rad, Germany). Proteins were transferred to polyvinylidene difluoride (PVDF) membranes (Biorad, Germany) using a Trans-Blot Turbo Blotting System (Bio-Rad, Germany). After this, an antigen-blocking step (5% non-fat milk powder, 1X Tris-buffered saline, 1% Tween-20) was performed, and the PVDF membrane was then incubated overnight at 4°C with primary antibody followed by secondary antibodies (see Supplementary information) for 1 hour. The antibodies were detected using enhanced chemiluminescence (ECL) (Bio-Rad, Germany), and the signals were detected by an INTAS Chemocam Imager (Royal Biotech GmbH, Germany). Immunoreactive bands were quantified by using the ImageJ software (http://rsb.info.nih.gov/ij/index.html) and the strongest band at ∼40 kDa of a Ponceau S stain of the membrane was used for normalization [18]. Please see the Supporting Information for detailed experimental procedures and included antibodies.

### Metabolomics and GC-MS analysis

For untargeted metabolomics, differentiated muscle cells and plantaris muscle tissue were prepared based on [19], and a Nexera UHPLC system (Shimadzu, Nakagyo-ki, Kyoto, Japan) coupled with a Q-TOF mass spectrometer (TripleTOF 6600, AB Sciex, Framingham, MA, USA) was used. Separation of the samples was performed using a HILIC UPLC BEH Amide 2.1 × 100, 1.7 μm analytic column (Waters Corp., Milford, MA, USA), and a reversed-phase Kinetex XB-C18, 2.1 × 100, 1.7-μm analytical column (Phenomenex, Torrance, CA, USA), respectively. The ^13^C labelling of the protein-bound alanine (product) was determined using gas chromatography-combustion-isotope ratio mass spectrometry (GC-C-IRMS) following derivatization as its MCME derivative. The media alanine (precursor) enrichment is determined by gas chromatography mass spectrometry. Unfortunately, we could not differentiate based on retention time, MS1 and fragmentation between glucose-6-phosphate, fructose-6-phosphate or glucose-1-phosphate which is why we took them together and referred to them as phosphorylated sugar. Please see the Supporting Information for detailed experimental procedures.

### Bioinformatics analysis of human data

Publicly available summary statistics [20] from the published genome-wide association study (GWAS) on appendicular lean mass of UK Biobank participants (n=450,243) [21, 22] were used to identify possible associations (p < 5.0e-8) between single nucleotide polymorphisms (SNPs) located in or near genes involved in the regulation of the serine synthesis pathway and its downstream enzymes and appendicular lean mass. In the next step, the Genotype-Tissue Expression (GTEx) portal [23] was used to analyze the association between the lean mass related SNPs and expression of specific genes in different tissues (p < 0.05). Please see the Supporting Information for detailed experimental procedures.

### Statistical Analysis

Data are expressed as mean ± standard error of means (SEM) for the indicated number of observations. Data were assessed for statistical significance by one-way or two-way ANOVA tests, followed by Tukey HSD, or by using a two-tailed Student’s t-test, as appropriate. A p-value of <0.05 was considered statistically significant. The reported unadjusted p values for all metabolites in the cell culture study were subsequently adjusted to account for multiple hypotheses testing. The false discovery rate (FDR) method of Benjamini and Hochberg was used to perform the adjustment (FDR<0.1). Unlike in vitro, we did not find significant results in the *in vivo* study after adjusting for multiple hypotheses testing FDR<0.1. We therefore took metabolites into account of the *in vivo* study, which were significant after adjusting for multiple hypotheses testing FDR<0.2 *in vivo*, and were also significant in the *in vitro* experiment. The specific statistical tests that were used are indicated in the figure legend. Please see the Supporting Information for detailed experimental procedures.

### Systematic Review

This systematic review is registered on PROSPERO (access number: CRD42022318998; (https://www.crd.york.ac.uk/PROSPERO/). One researcher searched in PubMed, Scopus and metabolite using the PICO framework leading to the following search strategy: ((mice or mouse) and (metabolomics or metabolome or metabolomes) and (skeletal muscle)) and ((increase or increased or increases or increasing or increasingly) or (mass gain) or (hypertrophy)) until May 11, 2022. Two researchers then screened the search results. Two researchers individually assessed the quality of each individual study using the 10-item checklist of CAMARADES (Collaborative Approach to Meta Analysis and Review of Animal Experimental Studies) (Table S2). A metabolite set enrichment analysis was performed using the MetaboAnalyst 5.0 web platform. Please see the Supporting Information for detailed experimental procedures.

## RESULTS

### Muscle growth stimulation with Insulin-like growth factor-1 increases the concentration of most of the glycolytic intermediates in C2C12 myotubes

To study the metabolic consequences of stimulation and inhibition of muscle protein synthesis in differentiated C2C12 muscle cells *in vitro*, we treated myotubes with 100 ng/mL of IGF-1 and/or 10 ng/mL rapamycin for 48 h. As expected, IGF-1 increased myotube diameter whereas rapamycin prevented the increase induced by IGF-1 (ANOVA, all p<0.001) (Supplementary Figure S1A). Treatment with IGF-1 also increased the phosphorylation of the mTORC1 downstream target 70-kDa ribosomal protein S6 kinase (p70S6K) after 2 hours (Figure S1B), which was accompanied 48 hours later by accretion of protein mass (Figure S1C) implying a positive protein balance, while rapamycin completely abolished basal and IGF-1 stimulated phosphorylation of p70S6K.

Next, we analyzed the metabolome of myotubes treated with vehicle control, IGF-1, or rapamycin for 48 h. We identified 155 metabolites from myotubes of which 28 showed significant differences between IGF-1 and control, 60 between IGF-1 and rapamycin, and 54 between control and rapamycin group, respectively (false discovery rate [FDR] <0.1; Figure 1A). Principal component analysis of the three treatment groups showed a clear separation of the three experimental groups (Figure 1B). IGF-1 treatment changed the concentration of several amino acids (Figure 1C) and glycolytic metabolites (Figure 2 & S2A).

**Figure 1.**
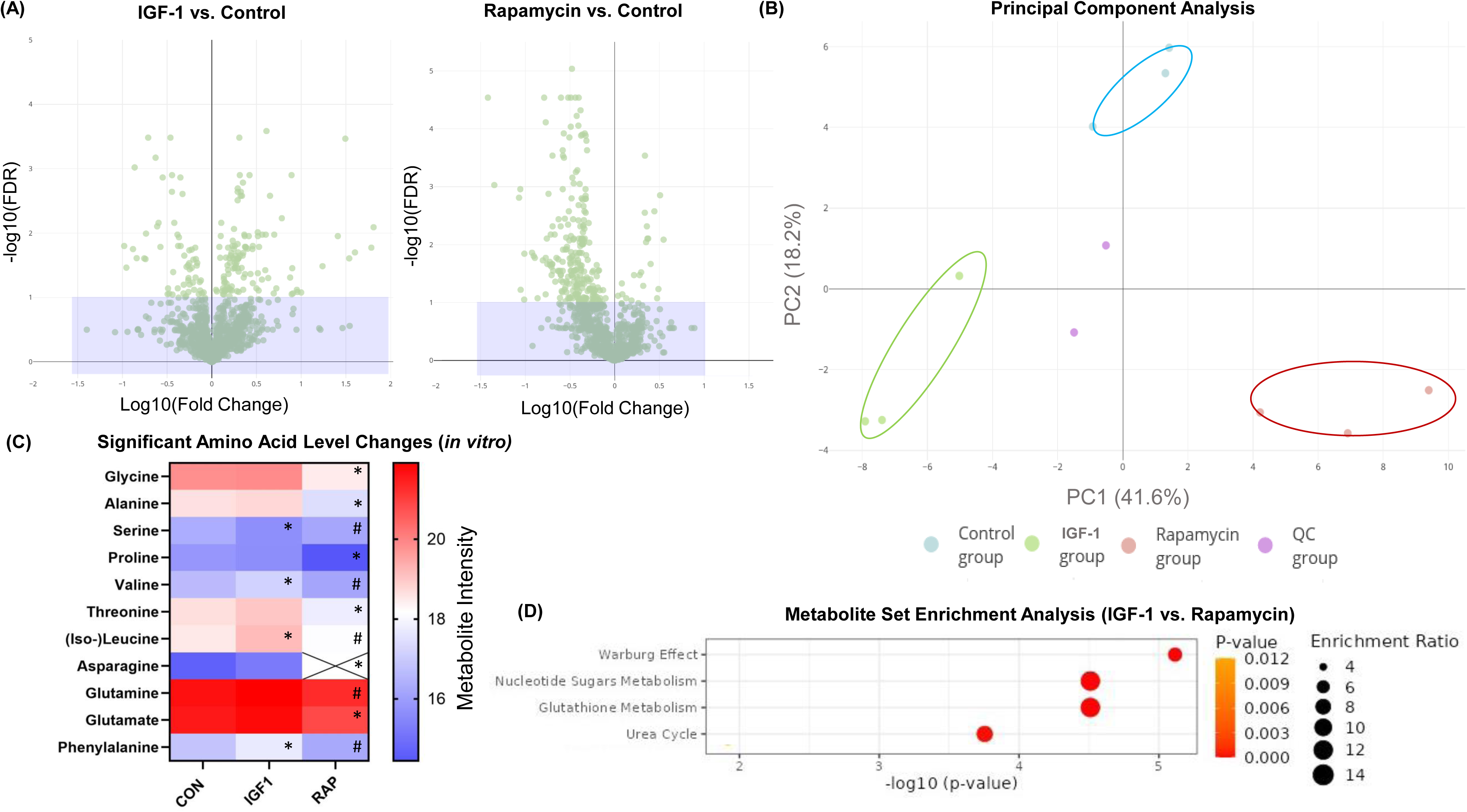
(A) Volcano plots displaying the false discovery rate (FDR)-values (−log10) versus log10 fold changes of all features (i.e. both unannotated and annotated metabolites) of LC-MS/MS measurements (negative ion mode) between Insulin-like growth factor-1 (IGF-1) and vehicle control (left) and between vehicle control and rapamycin (right) treatment in differentiated C2C12 muscle cells; features with FDR < 0.2 are above purple box. (B) Principal component analysis (PCA) of metabolites intensities of LC-MS/MS measurements (negative ion mode). (C) Heatmap of significant amino acid level changes after vehicle control, IGF-1 or rapamycin treatment in vitro. Crosses indicate that the metabolite is below the detection limit. *Significant differences between treatment (IGF-1 or Rapamycin) compared to vehicle control; # Significant differences between IGF-1 compared to Rapamycin, unpaired two-tailed Student’s t-test (FDR<0.1). (D) Enrichment analysis with MetaboAnalyst 5.0 between IGF-1 and rapamycin treatment. CON = Control; IGF1 = Insulin-like growth factor-1; RAP = Rapamycin.

**Figure 2.**
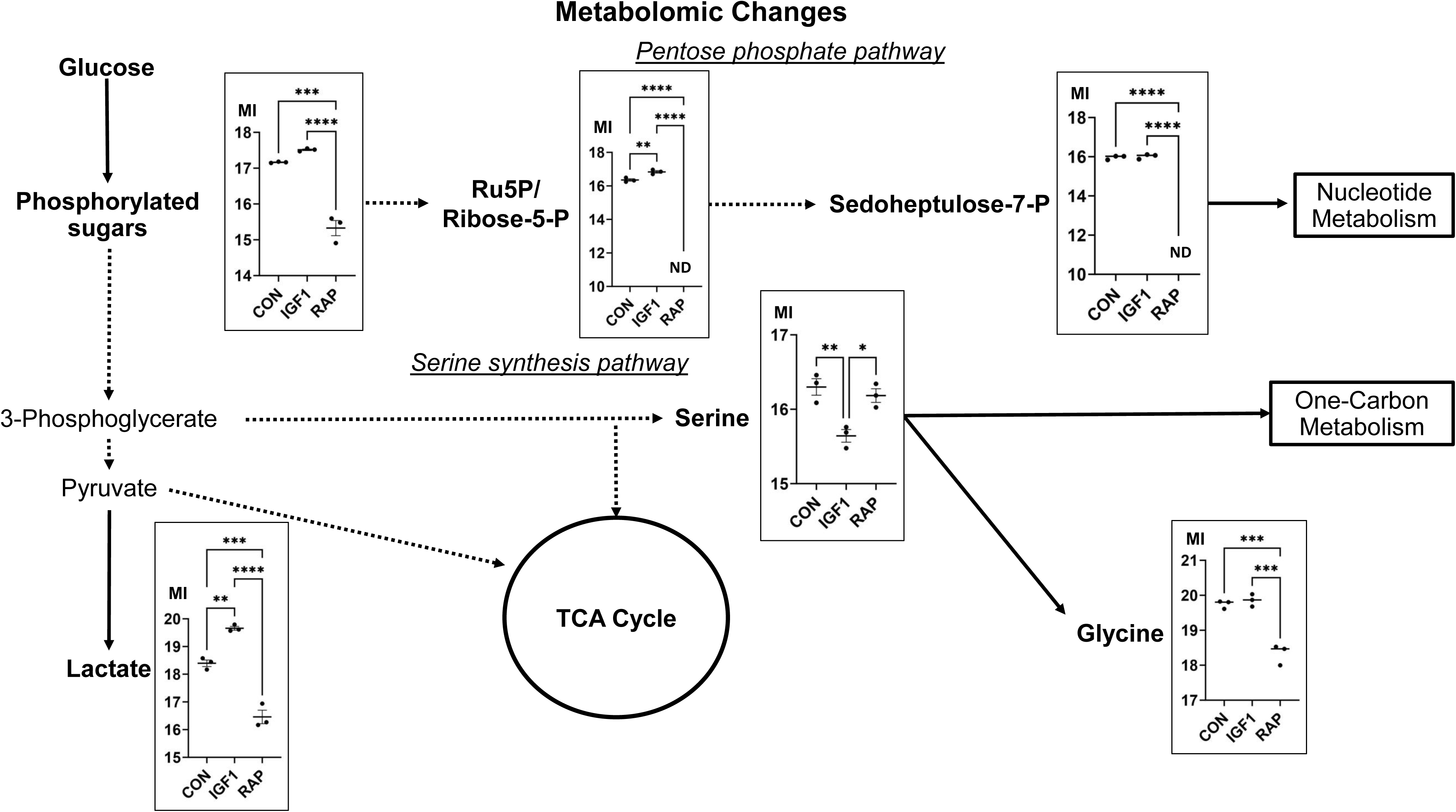
Metabolomic changes in differentiated myotubes treated with either vehicle control, Insulin-like growth factor-1 (IGF-1; 100 ng/mL) or rapamycin (10 ng/mL) for 48 h (n = 3). Metabolites highlighted in bold were detected by untargeted metabolomics. Frames show significant metabolite level changes between conditions (FDR<0.2). Metabolite intensity (MI) is represented on a Log2-scale. CON = Control; IGF1 = Insulin-like growth factor-1; ND = Not detected; P = Phosphate; RAP = Rapamycin TCA = Tricarboxylic acid cycle.

To identify differences in the metabolism between scenarios of robust skeletal muscle growth and its absence, we conducted a metabolic set enrichment analysis of distinct metabolite concentrations subsequent to the administration of IGF-1 when compared to rapamycin (Figure 1D), as well as of IGF-1 compared to control (Figure S2B), and rapamycin compared to control (Figure S2C). These analyses were consistent with the Warburg effect. We then analyzed the metabolic changes within the myotubes and focused on the metabolites that are associated with the glycolysis and to its linked anabolic pathways, which had altered (FDR<0.1) their concentration in response to IGF-1 or rapamycin treatment when compared to vehicle controls [Figure 2 and Supplementary table S3(C) and S4(C)]. IGF-1 treatment significantly increased the concentration of lactate by 139% in myotubes (p<0.001). With rapamycin treatment, the concentration of lactate decreased by 73.3% (p=0.002). An elevation of lactate is caused by increased glycolysis that is a phenomenon of the Warburg effect predominantly observed in cancer cells [5]. Concomitant with these changes, the concentration of phosphorylated sugar (the detection method used could not differentiate between glucose-6-phosphate, fructose-6-phosphate or glucose-1-phosphate) was higher by +27.5% (p<0.001) and of ribose-5-phosphate/ribulose-5-phosphate of the pentose phosphate pathway by +40% (p=0.01) after IGF-1 treatment. Phosphorylated sugar was reduced by 71.4% and ribose-5-phosphate/ribulose-5-phosphate as well as sedoheptulose-7-phosphate became undetectably low after rapamycin treatment (all p<0.001; Figure 2).

Of the 11 amino acids detected, IGF-1 treatment elevated the concentration of three (leucine/isoleucine, phenylalanine, and valine) amino acids, and rapamycin reduced the concentration of six (alanine, asparagine, glycine, glutamate, proline, and threonine) when compared to vehicle control (all p<0.01; Figure 1C). Of the detected amino acids, only serine had a lower concentration in IGF-1 treated myotubes compared to vehicle control (-36.8%; p=0.009). C2C12 myotubes consume rather than release serine into the media [24]. Thus, a decrease in serine concentration in the myotubes can either be based on a decreased activity of serine-producing enzymes or an elevated activity of serine-consuming enzymes [25]. The concentration of glycine (synthesized downstream of serine) did not change with IGF-1 treatment and the concentration decreased by 64% when treated with rapamycin (p=0.001). Thus, we assume that decreased serine concentration after IGF-1 treatment may indicate a high flux through the serine synthesis pathway due to an elevated demand of the glycine cleavage system, which refuels one-carbon metabolism [26]. Further, IGF-1 treatment increased metabolites linked to pyrimidine metabolism, i.e., uridine 5’-monophosphate (+82%; p=0.003) and uridine 5’-diphosphate (+130%; p=0.004) compared to control treatment, whereas the former was decreased by 58.7% in rapamycin group, when compared with vehicle control (p=0.016). The pyrimidine biosynthesis pathway is necessary for fundamental cellular functions such as for the DNA and RNA biosynthesis as well as for glucose metabolism.

Unlike glycolytic intermediates, TCA cycle metabolite concentrations changed little after IGF-1 treatment (Supplementary table S3-4). However, the intermediates fumarate and malate were decreased by ∼70% in myotubes treated with rapamycin (both p<0.0025; Figure S2A). Further, IGF-1 treatment increased the oxidized form of glutathione 10-fold indicating that IGF-1-induced muscle hypertrophy may elevate oxidative stress in myotubes.

A small fraction of glucose from glycolysis can also be shuttled into the hexosamine biosynthetic pathway that generates uridine diphosphate (UDP)-sugar precursors. These activated sugars are required for an often overlooked post-translational modification called *O-* linked glycosylation [27, 28]. The metabolomics analysis also revealed changes of metabolites of the hexosamine biosynthetic pathway (Figure S3A). Hypertrophy-stimulation by IGF-1 treatment increased the concentration of nucleotide sugars including UDP-N-acetylgalactosamine (UDP-GalNAc) (+76%), and UDP-glucuronic acid (+262%; both p<0.005). By contrast, rapamycin treatment decreased concentration of UDP-glucuronic acid (-73.3%; p=0.007), and glucosamine-6-phosphate as well as UDP-galactose concentrations to levels under the detection limit.

To investigate whether the changes of metabolite concentrations were accompanied by changes in the expression of enzymes that synthesize or metabolize these metabolites, we measured the expression of genes that encode metabolic enzymes. Elevated gene expression of key enzymes of the glycolysis and its parallel biomass pathways after treatment with IGF-I further supported the metabolic rewiring during skeletal muscle hypertrophy (Figure 3A). More specifically, glucose-6-phosphate dehydrogenase X (*G6pdx*), which encodes the rate-limiting enzyme of the oxidative pentose phosphate pathway, increased 3-fold in response to IGF-1 treatment. The serine-pathway-related enzymes *Phgdh* and mitochondrial serine hydroxymethyltransferase-2 (*Shmt2)* also changed between the intervention groups, and especially *Shmt2* increased ∼1.7-fold in abundance in hypertrophying muscle cells. The upregulation of these enzymes further suggests that the lower concentration of the serine metabolite as mentioned above was indeed caused by an increased flux through the serine synthesis pathway. Given the significance of metabolites of the hexosamine pathway in our *in vitro* analysis, we also assessed the gene expression of O-glycosylation-related enzymes and the abundance of total amount of glycosylated proteins in myotubes. However, we did not observe any significant differences between the groups at either the protein (Figure S3B) or gene level (Figure S3C).

**Figure 3.**
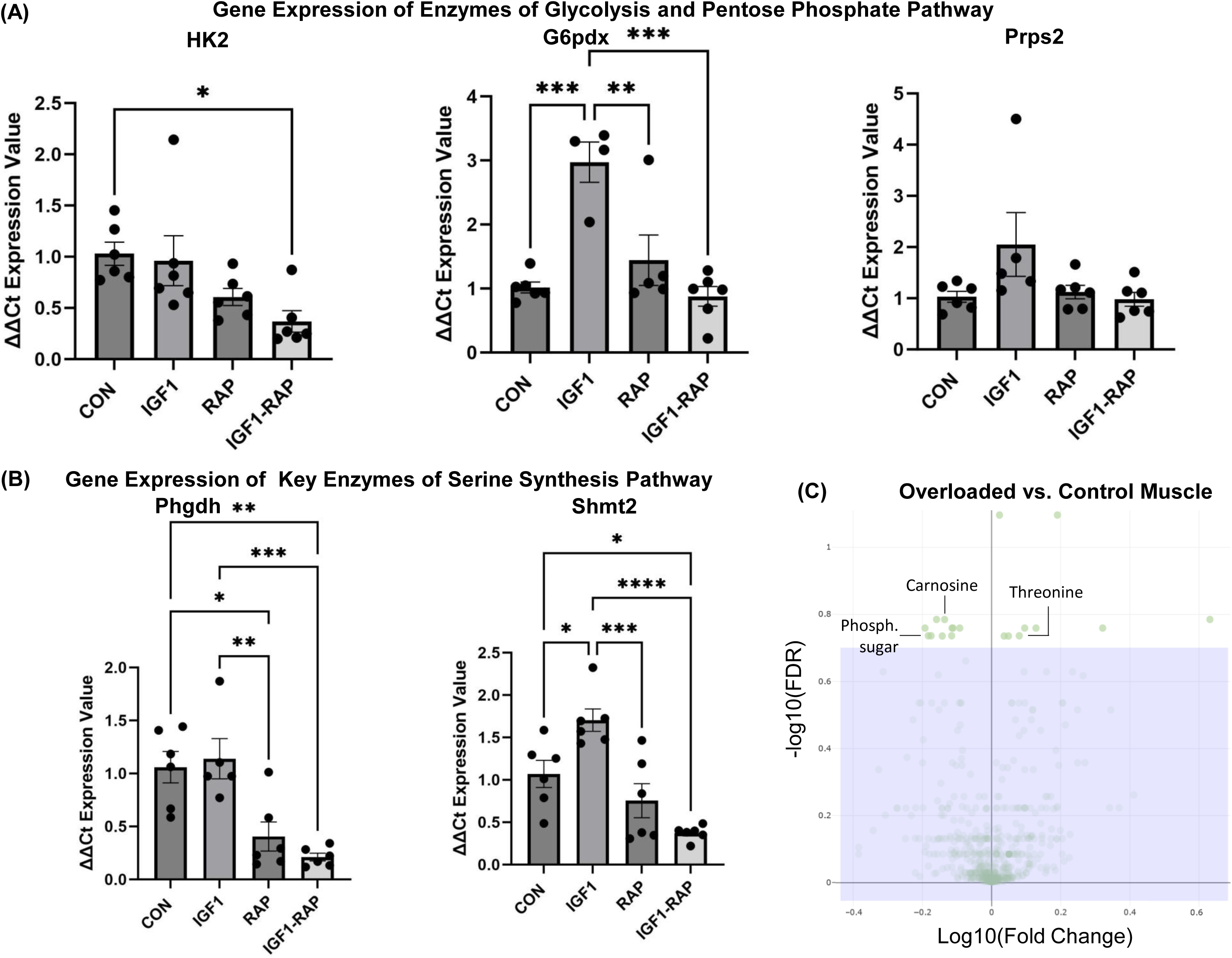
(A) Gene expressions of enzymes of glycolysis and pentose phosphate pathway including hexokinase 2 (Hk2), glucose-6-phosphate 1-dehydrogenase X (G6pdx) and phosphoribosyl pyrophosphate synthetase 2 (Prps2). (B) Gene expressions of key enzymes of the serine synthesis pathway - phosphoglycerate dehydrogenase (Phgdh) and serine hydroxymethyltransferase-2 (Shmt2). (C) Volcano plots displaying the false discovery rate (FDR)-values (−log10) versus log10 fold changes of all features between overloaded and control mice plantaris muscles (negative ion mode); features with FDR < 0.2 are above purple box. CON = Control; IGF-1 = Insulin-like growth factor-1; Phosph. sugar = phosphorylated sugar; RAP = Rapamycin. *Significant differences between indicated conditions, one-way ANOVA with Tukey HSD post-hoc test (p<0.05). Data are expressed as mean ± standard error (SEM).

To summarize, the data presented suggest that treating myotubes with IGF-1 results in a metabolic reprogramming that enhances the use of glycolytic intermediates for anabolic pathways. Conversely, rapamycin decreases myotube anabolism, and this is accompanied by lower concentrations of metabolites involved in anabolic pathways.

### Concentration of phosphorylated sugar also changes in hypertrophying murine muscle *in vivo*

To study the muscle metabolome response to a hypertrophic stimulus *in vivo*, we used an established muscle overload model [29]. Here, we induced compensatory hypertrophy of the right plantaris muscle in eight-month-old mice (n=6; body weight=31.3±2.6 g) by denervation of gastrocnemius and soleus muscles. The advantage of this model is that the plantaris muscle exhibits significant physiological hypertrophy by about 32% in 6 weeks [30], and yet is associated with less invasive surgery and inflammatory response than during overload induced by ablation of synergists. We chose to examine the muscle metabolome at 14 days after denervation at a time when there was yet only a small 13.5 ± 3.5% (p=0.007) increase in skeletal muscle mass, when compared to intact contralateral plantaris muscle, to capture a timepoint of a physiological and stable active growth in muscle size.

After rapid quenching of the muscles, we determined metabolite concentrations using an untargeted LC-MS/MS method (FDR <0.2; Figure 3C). Two metabolites exhibited alterations both in hypertrophied muscles in mice *in vivo* as well as in hypertrophying myotubes *in vitro.* Similar to the hypertrophy-stimulation with IGF-1 *in vitro*, the concentration of the dipeptide L-carnosine (known for its antioxidant activity and antiglycation effects) was decreased by 26.6% (p=0.001) during skeletal muscle growth *in vivo* (Supplementary table S5-6). However, phosphorylated sugar concentration showed contrasting trends, decreasing by 32.2% (p=0.004) in the overloaded muscle *in vivo*, while increasing in the IGF-1 stimulated myotubes *in vitro*, as mentioned before. When comparing the results of the *in vivo* muscle overload model with metabolite concentration changes between hypertrophy stimulating IGF-1 group and the absence of muscle growth represented by the rapamycin group *in vitro*, the concentration of threonine (*in vivo* +24.2%; p=0.009) was increased in both experimental models of muscle hypertrophy. We also tested the total muscle O-N-acetylglucosamine (*O*-GlcNAc)-modified proteins in the *in vivo* overloaded plantaris muscles and could not find any differences compared to the control muscles (p>0.05; Figure S3D).

### Systematic review further demonstrates a metabolic rewiring of glycolytic intermediates in growing muscles

Currently, no single metabolomics methodology covers the complete collection of metabolites in tissue [31]. To address this issue, we compared our findings to previously published metabolomics datasets of *in vitro* murine muscle cells and *in vivo* murine experiments to gain a comprehensive insight in metabolomic changes during and after skeletal muscle hypertrophy. For this, we performed a systematic review (PROSPERO access number: CRD42022318998) following PRISMA guidelines [32] and searched the literature in three literature databases (Pubmed, Scopus, Metabolight) using the PICO framework. To assess the quality and risk of bias of the *in vivo* studies, the ‘Collaborative Approach to Meta-Analysis and Review of Animal Data from Experimental Studies’ (CAMARADES) was used (Table S2). The results display metabolites that showed significantly changed concentrations in at least two experiments (Figure S4).

**Table 1.** Design and characteristics of the individual studies (n=8) included in the systematic review.

| Author (year) | Model Intervention (n) | Model Control (n) | Sex | Body Weight | Tissue analysed | Muscle weight | Age [weeks] | Intervention | Detection method |
| --- | --- | --- | --- | --- | --- | --- | --- | --- | --- |
| Baumert et al. (2022) | C2C12 muscle cells | C2C12 muscle cells | NA | NA | NA | NA | 5d after different. | IGF-1 incubation | LC-MS/MS |
| Baumert et al. (2022) | C57BL/6 mouse (6) | C57BL/6 mouse (6) | Male | 31.3g | Plan | C: 21mg<br>I: 24mg | 32 | Synergist ablation | LC-MS/MS |
| Cheng et al. (2014) | Akt1 DTG mouse (6) | C57BL/6 mouse (6) | Male | C: 38g<br>I: 38g | GA | C: 180mg<br>I: 220mg | 12 | Akt1 induction | H NMR |
| Collins-Hooper et al. (2015) | Mstn<sup>-/-</sup> mouse (Nk) | C57BL/6 mouse (Nk) | Both | C: 26g<br>I: 22g | GA | C: 100mg<br>I: 180mg | 16 | Myostatin KO | H NMR |
| Hoshino et al. (2020) | C2C12 muscle cells | C2C12 muscle cells | NA | NA | NA | NA | 6d after different. | EPS | CE-TOF-MS |
| Jiang et al. (2021) | C57BL/6 mouse (16) | C57BL/6 mouse (16) | Both | C: 26g<br>I: 26g | GA | C: 0.42%*<br>I: 0.50%* | 8 | Body Vibration | LC-MS/MS |
| Weyrauch et al. (2020) | C57BL/6 mouse (7-9) | C57BL/6 mouse (7-9) | Both | 27g | Plan | C: 21mg<br>I: 30mg | 6 | Synergist ablation | UPLC-MRM/MS |
| Wu et al. (2017) | Akt1 DTG mouse (Nk) | C57BL/6 mouse (Nk) | Both | C: Nk<br>I: Nk | GA | C: 0.5%*<br>I: 0.7%* | 16 | Akt1 induction | LC-MS/MS |

After removing duplicates, we screened 269 articles. Next to both present hypertrophying C2C12 muscle cells *in vitro* and plantaris muscle overload *in vivo* studies, we finally included another six publications that matched our eligibility criteria (see method section), which investigated changes in the metabolome of skeletal muscle tissue *in vivo* or myotubes *in vitro* during skeletal muscle hypertrophy [13, 14, 33–36]. We found 22 metabolites that were identified in at least two experiments (Figure 4A). About 73% of the observed metabolites that demonstrated changes in at least two experiments were detected in both study designs either analyzed hypertrophying mice muscles *in vivo* or in growing differentiated muscle cells *in vitro* (Figure 4B). This confirms that *in vitro* studies mimic the metabolic response to muscle hypertrophy stimulation. These results align to gene expression outcomes, which also revealed a great overlap between bioengineered muscles compared to acute resistance exercise in humans [37].

**Figure 4.**
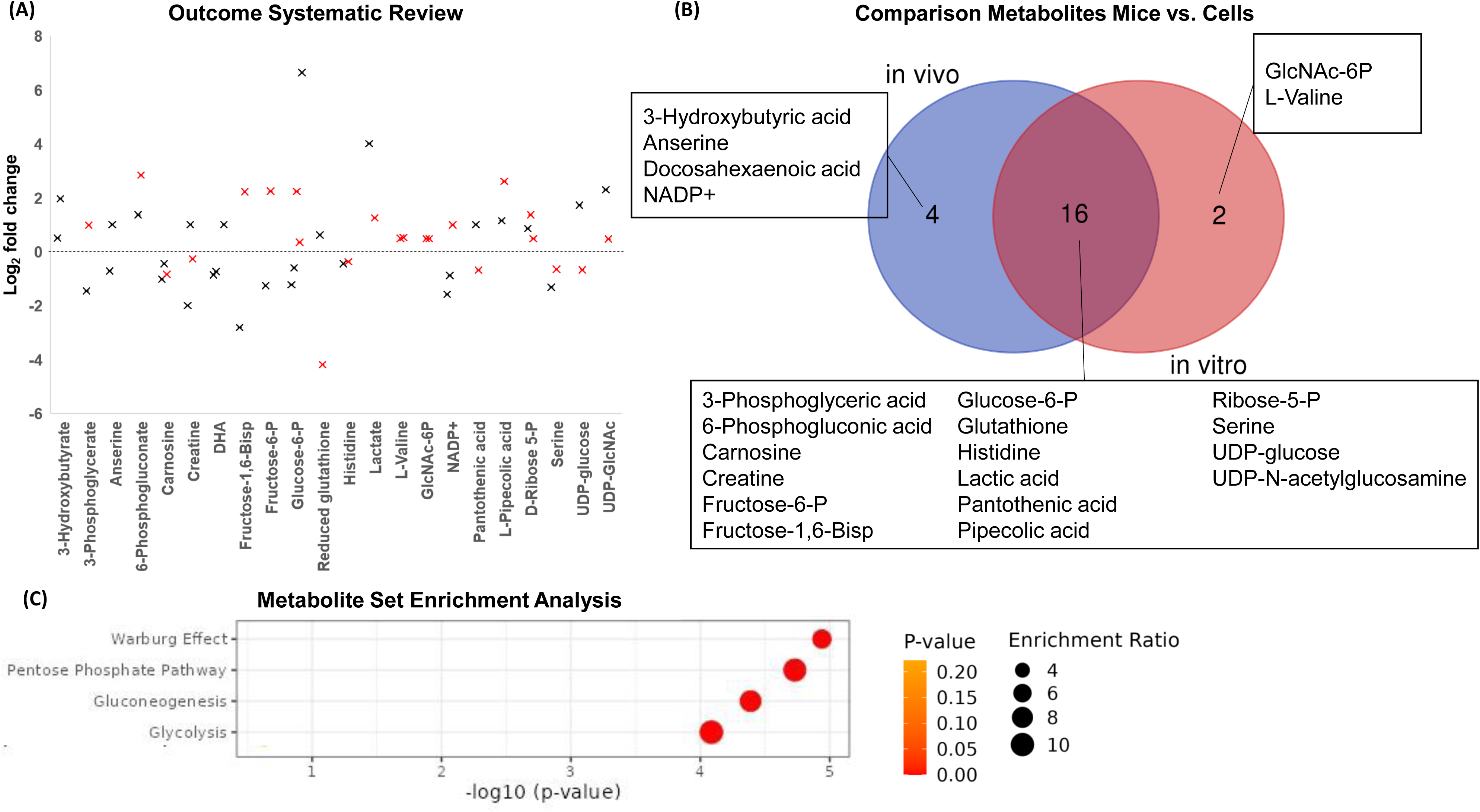
(A) Metabolic changes in response to skeletal muscle hypertrophy stimulation (log2 fold change versus rest), black cross – in vivo studies; red cross – in vitro studies. (B) Venn diagram showing overlap and unique metabolites associated with muscle growth between studies performed in mice in vivo (animals) or in vitro (cell culture). (C) Enrichment analysis with MetaboAnalyst 5.0. DHA = Docosahexaenoic acid; GlcNAc-6P = N-acetylglucosamine-6-phosphate; NADP+ = oxidized nicotinamide adenine dinucleotide phosphate; P = Phosphate; UDP = Uridine diphosphate.

Ten of these 22 metabolites were directly related to glycolysis and to its linked anabolic pathways. Generally, the concentrations of all included hexose phosphates (glucose-6-phosphate, fructose-6-phosphate, and fructose-1,6-bisphosphate) were elevated in cell culture studies *in vitro*. However, the prevailing trend was a decrease in these concentrations in *in vivo* hypertrophying mice muscles. Ribose-5-phosphate/ribulose-5-phosphate was also increased 1.5-2.5-fold. Both lactate and valine concentrations were increased up to 16-fold and 1.5-fold during muscle hypertrophy, respectively. Supporting the *in vitro* investigation presented here, serine concentration was decreased also in Cheng et al. [14] (Figure 4A).

Metabolic set enrichment analyses of different metabolite concentrations following muscle hypertrophy-inducing treatments revealed a significant association in the concentrations of metabolites linked to the pentose phosphate pathway, glycolysis and gluconeogenesis, all characteristic of the Warburg effect (Figure 4C). These results, similar to the observed changes in our muscle cell *in vitro* and mice study *in vivo*, further suggest that skeletal muscle hypertrophy, like in cancer, causes a metabolic reprogramming to generate biomass.

### [U-^13^C_6_]Glucose tracer confirms increased amino acid carbon incorporation in IGF-1 stimulated C2C12 myotubes

An observation that we made in our initial studies was a shift in the concentration of glycolytic intermediates and their associated anabolic pathways (e.g. the serine synthesis pathway) during muscle hypertrophy. This observation was supported by the metabolic set enrichment analysis of the systematic review that revealed an association with the Warburg effect. We next investigated the fate of glucose during skeletal muscle growth (Figure 5A). For this, we used stable isotope [U-^13^C_6_]glucose tracer to identify whether stimulation with IGF-1 alters the incorporation of glucose carbon into amino acids and, ultimately, into the protein biomass of C2C12 myotubes over 24 h. Intermediary metabolism of U-^13^C_6_ labelled glucose results in ^13^C labelled carbon being incorporated into non-essential amino acids. The enrichment of ^13^C-glucose within the conditioned media, measured in atom percent excess (∼23.5 APE), did not differ between the groups at 24 h post-tracer treatment (p=0.95; Figure 5A). However, the IGF-1 treated cells showed significant additional ^13^C-enrichment in the media lactate and alanine compared to control, suggesting that lactate and alanine was increasingly synthesized from glucose and then released into the media (Figure 5A). Glycine (p=0.068) and three distinct isotopologues of proline were detected, reflecting the different routes of carbon into *de novo* proline synthesis; Figure 5B) were also enriched with ^13^C in the IGF-1 group within the conditioned media, respectively, reflecting the flux of glucose carbon into the non-essential amino acid pools. The enrichments were slightly higher in the IGF-1 group, possibly reflecting fluxes through de novo synthesis pathways but also inhibition of breakdown. We did not observe any ^13^C-enrichment of serine in either the conditioned media or skeletal muscle protein. The limited ^13^C-enrichment of serine in the media indicates that differentiated skeletal muscle cells take up more serine than they release. It also suggests that glucose-synthesized serine is rapidly utilized in downstream pathways. To ascertain whether hypertrophic muscles consume serine rather than release it, we quantified the total serine metabolite concentration in the media. Our results imply that IGF-1-treated cells potentially consume more serine from the media compared to the control cells (Figure 5C). We then estimated the fractional rate of protein synthesis (FSR) with the ^13^C-enriched alanine that is incorporated into muscle protein, and FSR was elevated by ∼13% with IGF-1 treatment (p=0.001; Figure 5D). This indicates that ^13^C labelled carbon of glucose is increasingly converted to pyruvate, which is then transaminated to form alanine, which is then incorporated into protein.

**Figure 5.**
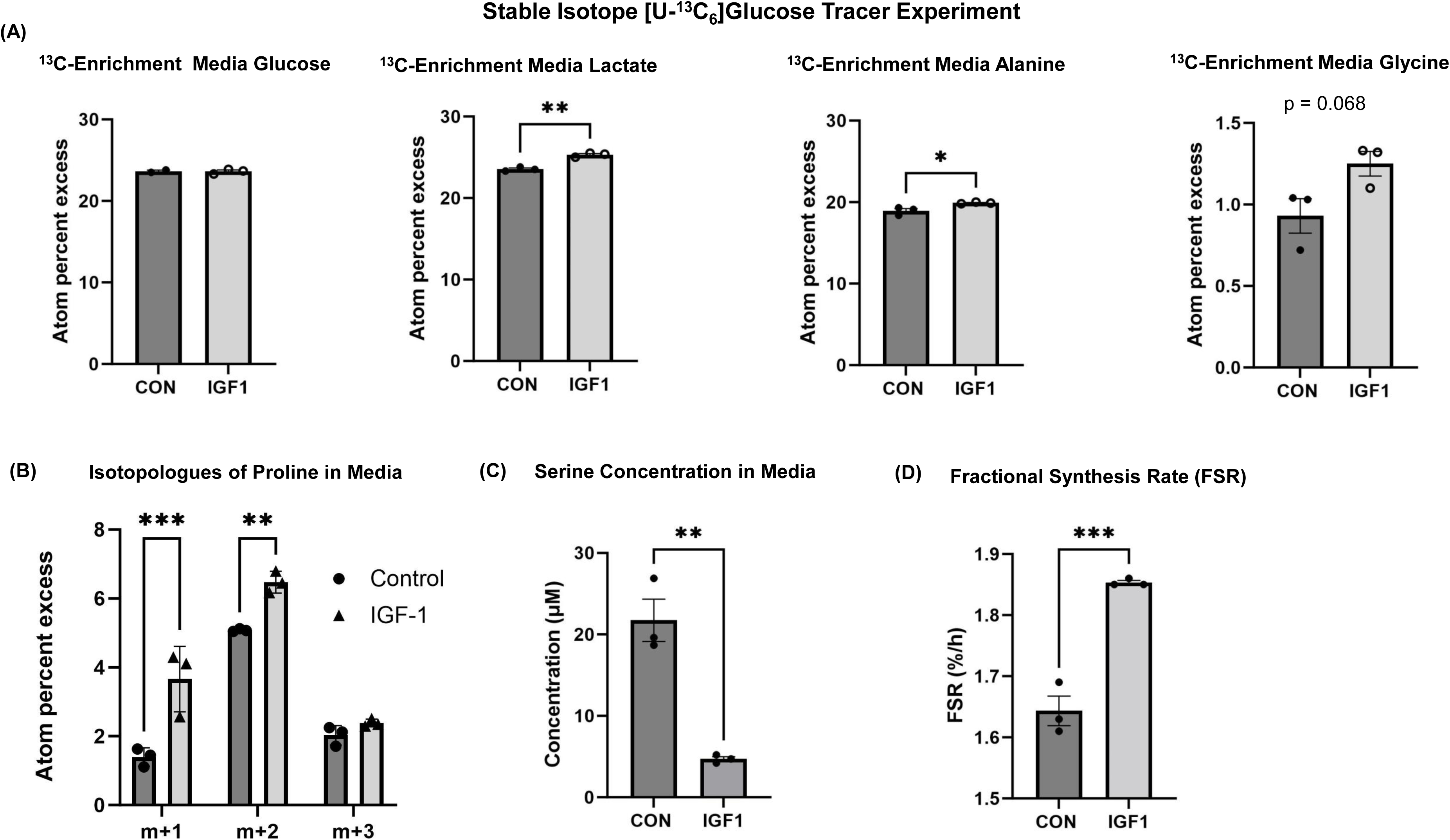
(A) ^13^C-Enrichment of glucose, lactate, alanine and glycine between IGF-1 and vehicle control in conditioned media after 24 h. (B) Isotopologues of Proline in Media; (C) Serine Concentration in conditioned Media (D) Fractional synthesis rate (FSR) based upon ^13^C-alanine incorporation; CON = Control; IGF-1 = Insulin-like growth factor-1; RAP = Rapamycin. *Significant differences between groups, unpaired two-tailed Student’s t-test (p<0.05). Data are expressed as mean ± standard error (SEM).

### Inhibition of the key serine synthesis pathway enzyme PHGDH reveal a key regulatory or allowing role of the serine synthesis pathway in muscle protein synthesis

Our metabolomics *in vitro* study as well as the systematic review revealed a decreased serine concentration in growing muscles and an upregulation of enzymes that are linked to the serine synthesis pathway. With stable isotope [U-^13^C_6_]glucose tracer, we also confirmed increased incorporation of glucose-derived carbon into glycine suggesting that there may be increased flux through this pathway during muscle growth. Further, we recently reported that Phgdh knock down is linked to a smaller myotube size [16]. Thus, we next inhibited Phgdh to investigate its role in muscle protein synthesis during muscle cell differentiation. Chemical inhibition of Phgdh with NCT-503 (50 µM) led to reduced protein synthesis by 58.1% (p<0.001) measured by puromycin labelling (Figure 6A), which resulted in a 36.8% lower total muscle cell protein concentration (Figure 6B). Extracellular lactate dehydrogenase (LDH) activity was not changed thus indicating that the Phgdh inhibition did not cause major cytotoxic effects (Figure 6C).

**Figure 6.**
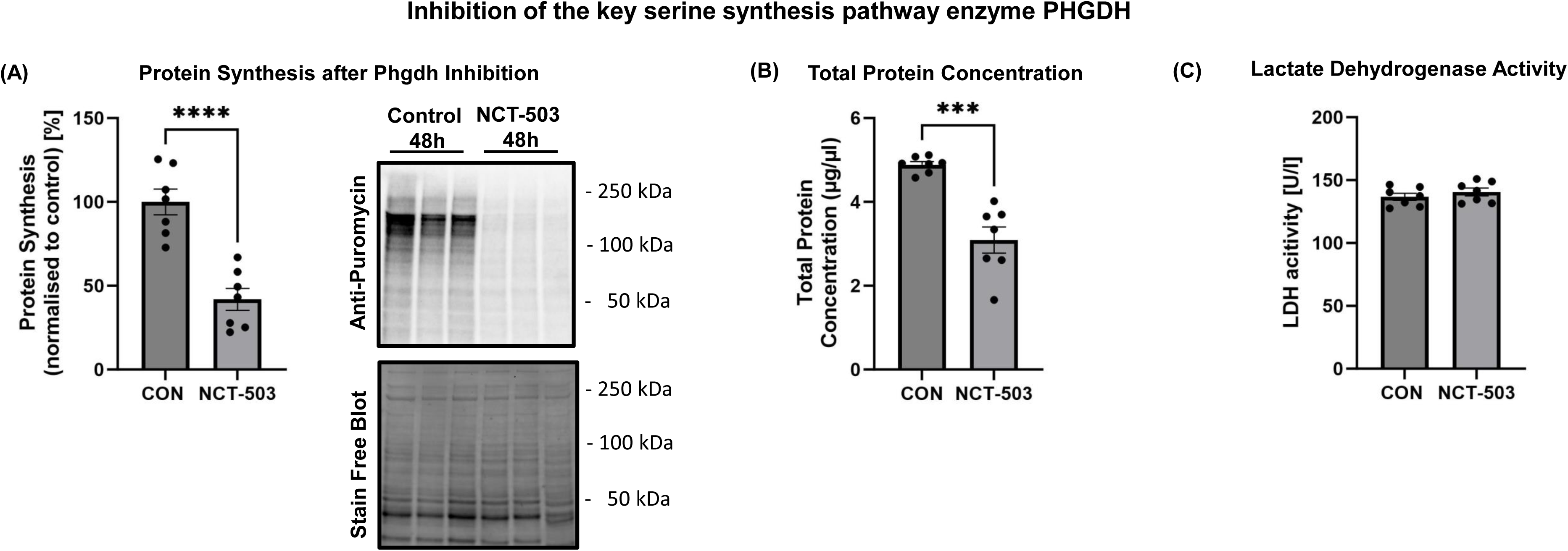
(A) Protein synthesis determined using puromycin, (B) total protein concentration of C2C12 myotubes, and (C) lactate dehydrogenase (LDH) activity in conditioned media measured including NCT-503 inhibitor or vehicle control (CON) after 48 h. *Significant differences between groups, unpaired two-tailed Student’s t-test (p<0.05). Data are expressed as mean ± standard error (SEM).

Collectively, these data suggest that the serine synthesis, pentose phosphate and hexosamine pathways are reprogrammed during skeletal muscle hypertrophy to support biomass production while inhibiting biomass pathways decreases protein synthesis. Skeletal muscle converts carbon from glucose into non-essential amino acids and the rate of incorporation increases during muscle hypertrophy.

### Variants of genes involved in serine synthesis pathway are associated with appendicular lean mass in the UK Biobank cohort

As we observed serine synthesis pathway enzyme to regulate cell anabolism and size, we next identified using publicly available datasets [20] three SNPs located in or near *PHGDH, PSPH* and *SHMT1* genes significantly associated with appendicular lean mass, which is a proxy variable for muscle mass, at the genome-wide significant level (p<5.0e-8) in the UK Biobank cohort (n=450,243). More specifically, we identified an association of the *PHGDH* gene rs12129705 A (p=1.2e-14, beta=0.0221), *PSPH* gene rs11982736 G (p=9.2e-17, beta=0.0217), and the *SHMT1* gene rs11867855 G (p=2.9e-9, beta=0.0132), which encodes the cytosolic form of SHMT, alleles with increased appendicular lean mass. Follow-up gene expression analysis using GTEx portal [23] revealed that these variants are functional and affect gene expression in various tissues, such as in skeletal muscle (*PSPH* rs11982736 gene variant; p=6.8e-74) and in testis (*PHGDH* rs12129705 and *SHMT1* rs11867855 gene variants; p<0.0000063).

## DISCUSSION

The major finding of this study suggests that a hypertrophying skeletal muscle remodels its metabolism so that carbon from glucose increasingly diverts into anabolic pathways for building biomass. This is indicated by altered intermediate metabolite concentrations of anabolic pathways. This was further supported by stable isotope [U-^13^C_6_]glucose tracer experiments that indicates that labelled carbon from glucose is incorporated into the non-essential amino acid alanine in normal growing myotubes and more labelled alanine is incorporated in hypertrophying myotubes. Further mechanistic analysis revealed that the inhibition of Phgdh suppresses muscle protein synthesis in growing myotubes. This finding, coupled with GWAS analysis on appendicular lean mass in UK Biobank participants, underscores the importance of the serine synthesis pathway in muscle maintenance and potentially in preventing muscle loss.

### Metabolic reprogramming

L-carnosine and phosphorylated sugar changed their concentrations in both hypertrophied mouse muscles *in vivo* and cultured hypertrophying myotubes *in vitro*. However, the levels of phosphorylated sugar (glucose-6-phosphate, fructose-6-phosphate or glucose-1-phosphate) exhibited contrasting trends: increasing in IGF-1 stimulated myotubes *in vitro* but decreasing in overloaded muscle *in vivo.* Our *systematic review* also showed that the concentration of hexose phosphates (glucose-6-phosphate, fructose-6-phosphate, and fructose-1,6-bisphosphate) were consistently elevated in studies using myotubes *in vitro*. In contrast, the majority of these concentrations exhibited a decrease in *in vivo* hypertrophying mice muscle experiments included in our review, with fructose-1,6-bisphosphate also showing a decrease after five-week-long resistance training regimen in healthy humans [38]. This suggests heightened metabolic activity during muscle growth, possibly due to increased demand for hexose phosphates as substrates for anabolic pathways. However, the availability of glucose and metabolites in cell culture media may not mirror *in vivo* conditions for (fasted) mice, prompting further exploration of dietary influences on muscle hypertrophy and metabolic reprogramming.

Of the detected amino acids, only the concentration of serine was lower in hypertrophy-simulated muscle cells. We also found that hypertrophy-stimulated muscles did not show any changes of Phgdh gene expression, the rate-limiting enzyme of the serine synthesis pathway, but had an increased expression of the *Shmt2* gene, that encodes an enzyme downstream of Phgdh. The gene encodes the serine hydroxymethyltransferase 2 enzyme that catalyzes the reversible reaction of serine to glycine, thus promoting the production of one-carbon units that are indispensable for cell growth [26]. Other studies also showed a reduction of muscle serine during muscle growth over time [14]] and that C2C12 myotubes actively uptake significant quantities of serine from the media and secretes glycine in the media [39]. Therefore, the lower serine concentration in hypertrophying muscles might be due to an increased activity of the serine consuming enzyme serine hydroxymethyltransferase 2 for providing precursors for macromolecules. In support of this, we found -58.1% lower protein synthesis in Phgdh-inhibited myotubes, demonstrating that blocking serine synthesis pathway limits protein synthesis. Further, the genetic variation data from the UK Biobank support the association between enzymes of the serine synthesis pathway (*PHGDH, PSPH*) and of *SHMT1* with the regulation of skeletal muscle mass. This extends beyond species-specific findings in mice, suggesting relevance of parallel pathways of glycolysis such as serine synthesis pathway on the regulation of muscle size at the population level.

We estimated with stable-isotope labelled [U-^13^C_6_]glucose tracers the fractional synthesis rate of ^13^C-enriched alanine that is incorporated into muscle protein and the results revealed an increase of 13% in the IGF-1 group compared to control. We also saw a small additional enrichment of carbon from glucose in the detected non-essential amino acids glycine, proline, alanine and especially lactate within the conditioned media, when myotubes were stimulated with IGF-1. This suggests that the rate of incorporation of amino acids into skeletal muscle is greater during muscle growth and that the increased carbon demand for the *de novo* biosynthesis is met by an elevated glycolysis combined with a redirection of glucose intermediates toward anabolic pathways.

In addition to serine and pentose phosphate pathways, glucose can also be directed towards the hexosamine biosynthetic pathway, resulting in the generation of UDP-sugar precursors. Our *in vitro* findings suggest that IGF-1 acts as an anabolic stimulus, while rapamycin serves as a catabolic stimulus, reprogramming these metabolic pathways. However, we did not observe a clear association with O-GlcNAcylation at either the genetic or protein level, in contrast to previous research [28]. The underlying reasons for this discrepancy remain unclear, suggesting that O-GlcNAcylation may not consistently change during muscle hypertrophy. Further mass spectrometry-based studies are needed to uncover more details of cellular glycosylation in the future.

Our investigation shed further light on the metabolism during muscle growth. Hypertrophying muscle rewires the metabolism accompanied with metabolic changes in pathways associated with the Warburg effect. The results of our investigation might help to develop effective nutritional and exercise therapies against muscle wasting during ageing and metabolic diseases. Future studies should investigate whether inhibition of other key enzymes parallel to glycolysis linked with the Warburg effect [10] will further shed light on the metabolic response to muscle hypertrophy. More critically, we underscore the necessity for further studies in human participants to corroborate and expand upon our findings, potentially paving the way for practical interventions in clinical settings.

### Limitations

We demonstrated a regulatory function of the serine synthesis pathway in muscle size regulation by inhibiting Phgdh with NCT-503. Further studies should use several inhibition and activation strategies *in vivo* for Phgdh and other key enzymes of the serine and pentose phosphate pathway. Additionally, our results of the increased Shmt2 gene expression and the relative lower metabolite concentration of serine in both muscle and conditioned media suggest an increased metabolite flux through the serine synthesis pathway. Our ^13^C-glucose analysis allowed us to detect ^13^C-enrichment in specific amino acids. To validate our findings, a comprehensive ^13^C-glucose metabolic flux analysis could directly assess the metabolic flux in the serine and other pathways. The ^13^C-enrichment in muscle protein alanine was increased. However, we have only measured FSR, and the enrichment could also reflect inhibition of muscle breakdown. Due to resource constraints, this study focused exclusively on male mice, potentially missing valuable insights into any sex-specific effects. This systematic review acknowledges a potential limitation in the comprehensiveness of its literature search. Despite our systematic and methodical approach, it is possible that not all relevant publications within the searched timeframe were identified and included. Further, we were unable to include the measurement of metabolite variation around its means due to a lack of information in some included publications so our analysis is not considered meta-analysis.

## CONCLUSION

Our results provide further evidence that muscle hypertrophy is associated with metabolic remodeling. Understanding of the mechanisms regulating skeletal muscle mass can assist in developing treatments for preventing and treating muscle weakness and atrophy.

## ACKNOWLEDGEMENT

We greatly acknowledge the support of Prof. Ken Smith (School of Medicine, University of Nottingham), who processed and analyzed the samples from the [U-^13^C_6_]glucose experiment, and of Dr Marius Garmhausen for his insightful feedback during the data analysis phase. The authors of this manuscript certify that they comply with the ethical guidelines for authorship and publishing in the Journal of Cachexia, Sarcopenia and Muscle [40].

## ETHICS STATEMENT

All procedures involving mice were approved by the Lithuanian State Food and Veterinary Service (Animal ethics number: 2018-06-14 Nr. G2-90) under consideration of the well-being of the animals and the 3R (replace, reduce, and refine) principle. The manuscript does not contain clinical studies or patient data.

## AUTHOR CONTRIBUTION

P.B., M.S., T.V., H.S., A.R., H.D., J.J.H. and H.W., conceived and designed the research; P.B., S.M., A.S., P.M., and M.D. performed the experiments; P.B., M.S., S.M., I.A., K.G., C.M., and M.G. analyzed the data; P.B., M.S., H.D., A.R., J.J.H., and H.W. interpreted the results of the experiments; P.B. prepared the figures; P.B. drafted the manuscript; P.B., M.S,, S.M., A.S., P.M., M.D., T.V., H.S., H.D., A.R., J.J.H., and H.W. edited and approved the final version of the manuscript.

## DATA SHARING

The datasets that support the findings of the animal study are available as Supplementary Information files. The human data presented in this study are publicly available online at https://genetics.opentargets.org (accessed on 20^th^ December 2023) and https://gtexportal.org/home/index.html (accessed on 20^th^ December 2023).

---

**Supplementary Figure S1.**
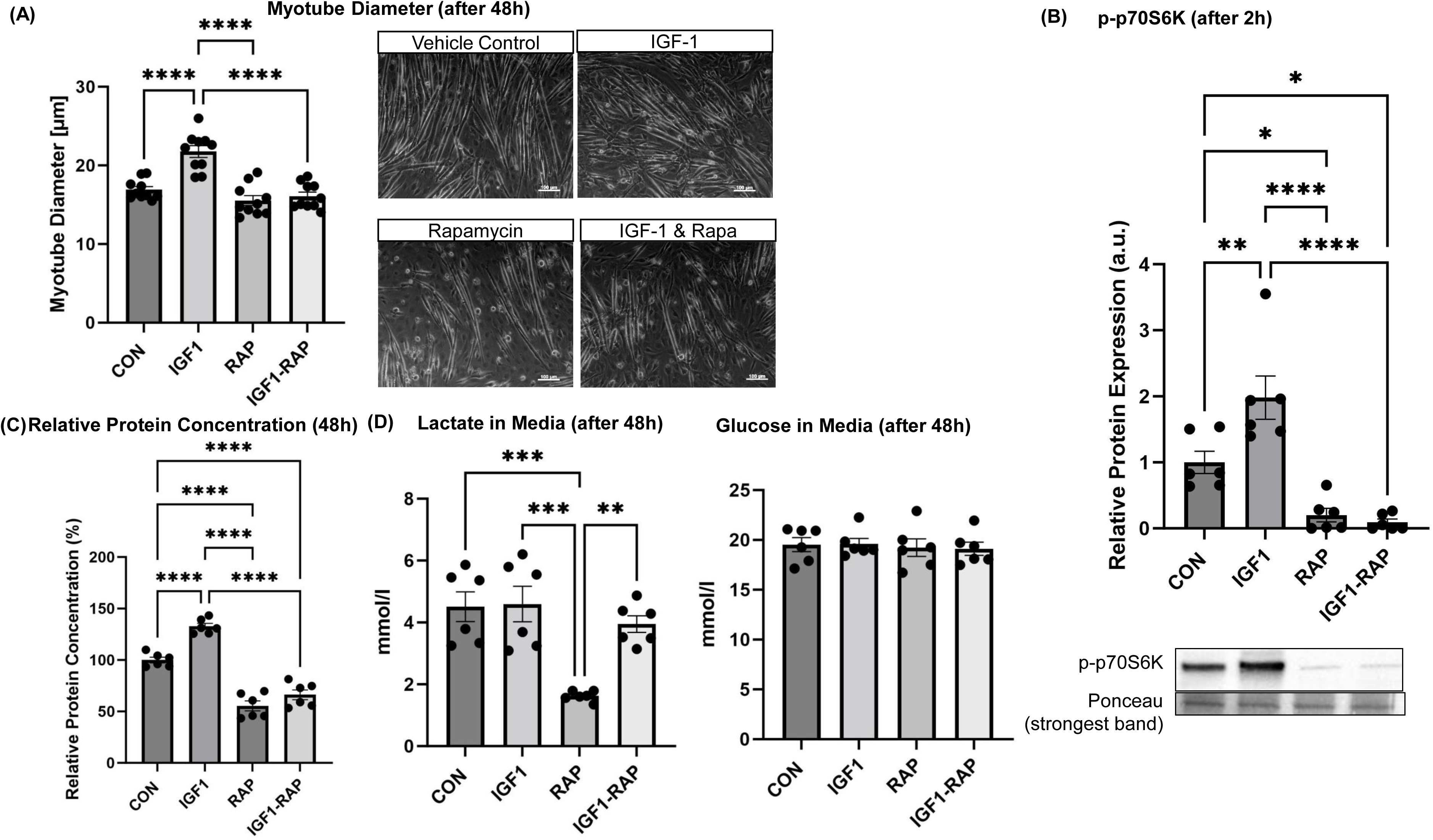

**Supplementary Figure S2.**
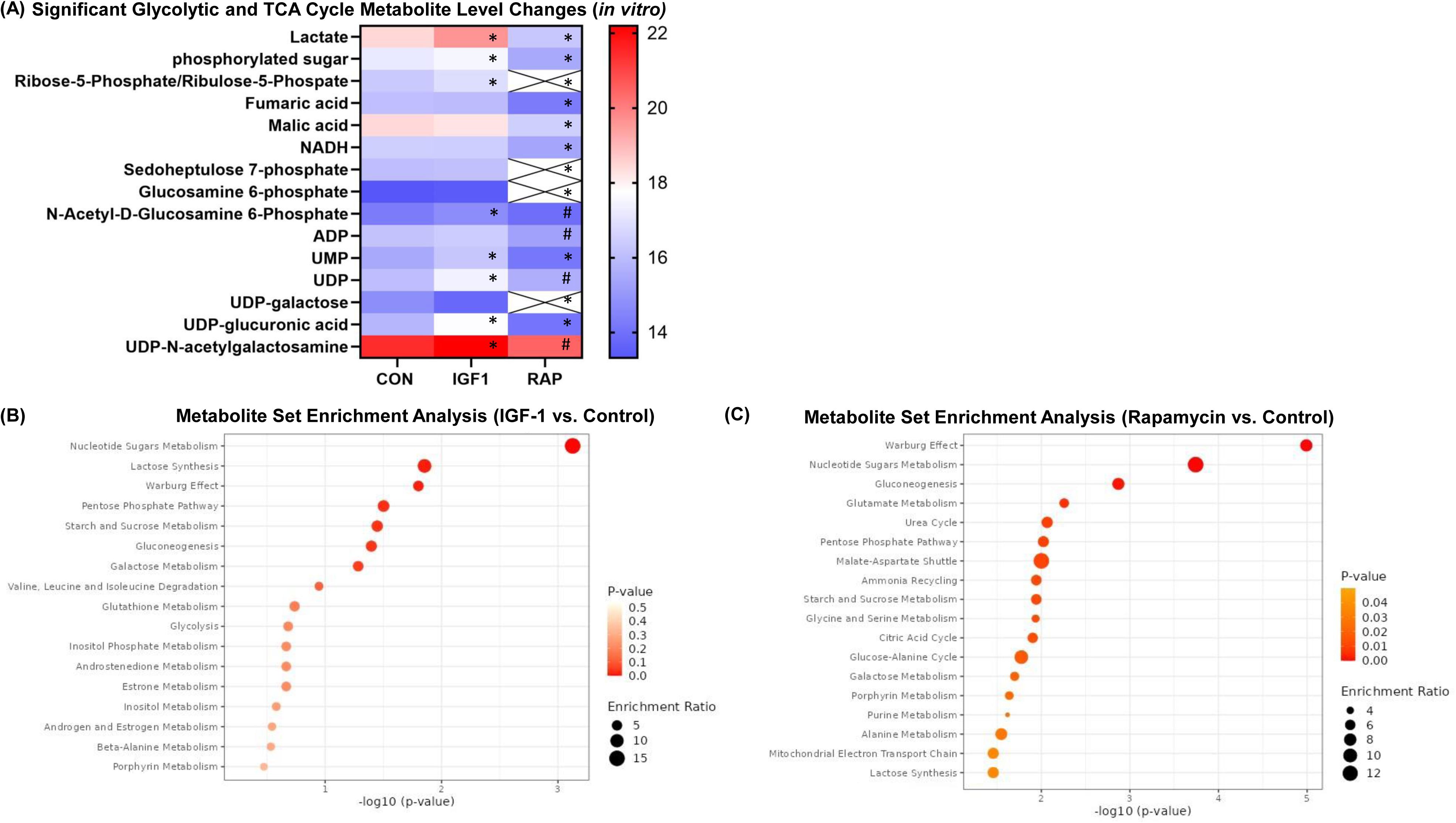

**Supplementary Figure S3.**
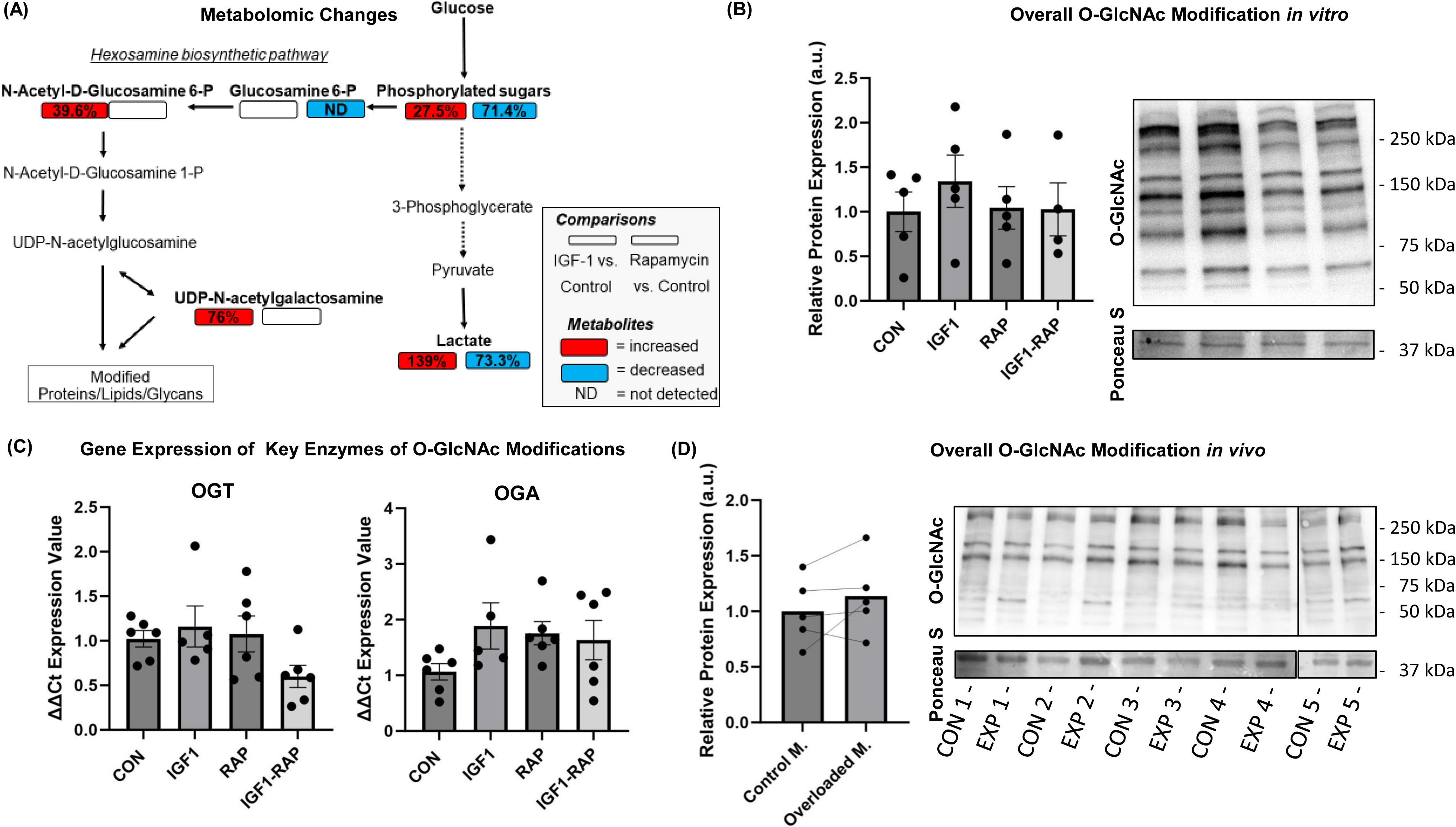

