## Supplementary Material for "Skeletal muscle hypertrophy rewires glucose metabolism: an experimental investigation and systematic review"

### MATERIAL AND METHODS

##### Animals and *in vivo* study design

All procedures involving mice were approved by the Lithuanian State Food and Veterinary Service (Animal ethics number: 2018-06-14 Nr. G2-90). Male C57BL/6J mice were used in all experiments. Mice were housed in standard cages, one to three mice per cage at a temperature of 22–24°C and 40–60% humidity. Animals were fed standard chow diet and received tap water ad libitum.

Six mice were subjected to muscle overload for 14 days starting at the age of 32 weeks, as described previously [1]. These animals were anesthetised with an intraperitoneal injection of ketamine (100 mg/kg, Ketalar, Pfizer, Belgium) and xylazine (10 mg/kg, Rompun, Bayer, Belgium), and acepromazine (3 mg/kg, Placivet, Kela, Belgium) administered intraperitoneally at a volume of 150 mL/25 g. After cessation of nociceptive responses, an incision was made in the popliteal area of the left hind limb to expose the tibial nerve with blunt dissection. The branches of the tibial nerve that innervate the soleus and gastrocnemius muscles were cut and small segments removed to prevent reinnervation. This strategy imposes an overload and subsequent compensatory hypertrophy of the plantaris muscle. The contralateral plantaris muscle of these mice served as internal control in analysis of muscle mass and metabolomics measurements. Mice were given buprenorphine (0.2 mg/kg) after surgery for pain relief once a day for 3 days and were monitored on a daily basis. After 14 days, mice were fasted overnight and they were sacrificed by the exposure to $CO_2$. Immediately afterwards, plantaris muscle was dissected, blotted dry, weighed to 0.1 mg using an analytical balance (Kern, ABS 80-4, Germany), frozen in isopentane (pre-cooled with liquid nitrogen), and stored at − 80 °C until further metabolomics analysis.

##### Cell culture and *in vitro* study design

Mouse C2C12 cells were a kind gift from Peter Meinke (Ludwig Maximilian University of Munich, Germany). Cells were tested at Eurofins Genomics forensic department in Ebersberg, Germany, and found negative for mycoplasma contamination. C2C12 myoblasts cells were cultured in Dulbecco's modified Eagle's medium (DMEM) (4.5 g/L Glucose; Gibco, Cat#11965-092, Waltham, MA, USA), supplemented with 10% foetal bovine serum (Sigma-Aldrich, F0804, NY, USA), and incubated at 37 °C in humidified air with 5% CO2. To induce differentiation, the myoblasts were cultured until confluence and then the medium was switched to DMEM with 0.2% FBS after washing the cells twice with Phosphate-buffered saline (PBS; Sigma-Aldrich). The proliferation and differentiation medium were refreshed every other day. After 72 h of differentiation, C2C12 Myotubes were serum starved (DMEM only) for 4 h and then treated with DMEM including 0.2% dialysed FBS (Thermo Scientific, Cat# A3382001, Gibco, USA) and vehicle control (10 mM HCL, 0.1% bovine serum albumin; BSA), IGF-1 (100 ng/ml; recombinant Human LONG $R^3$ IGF-1, Sigma-Aldrich, Cat# I1271, MO, USA), rapamycin (10 ng/ml; Calbiochem, Cat#553210, Watford, Hertfordshire, UK) and IGF-1 + rapamycin. for 48 h. For Phgdh inhibition experiments, we treated C2C12 cells with chemical PHGDH inhibitor NCT-503 (50 µM; MCE, HY-101966; MedChemExpress) five days after the initiation of differentiation for 48 h.

#### Myotube size measurement

Digital photos of cross-sections were captured using a Zeiss Axio Lab.A1 equipped with a digital camera (Zeiss, Germany) and Zeiss ZEN Software version 2.6 (Blue edition; Zeiss, Germany). Four photographs of each well were taken at 10x magnification after the 48-h treatment. The average diameter per myotube was evaluated at three points along the length of the myotube in a blinded fashion using ImageJ (http://rsbweb.nih.gov/ij/, National Institutes of Health, Bethesda, MD, USA; RRID:SCR_003070) and taking into account the pixel-to-aspect ratio. At least 50 myotubes per well were measured.

#### Conditioned Media Lactate dehydrogenase activity

Lactate dehydrogenase (LDH) enzyme activity (No. 981906, Thermo Fisher Scientific), a biomarker of cell rupture, was measured with an automated Indiko plus analyzer (Thermo Fisher Scientific) according to manufacturer's protocol from the media 48 h after the treatment with 50 µM NCT-503 or DMSO vehicle.

#### RNA isolation, reverse transcription, and quantitative real-time PCR

C2C12 myotubes were washed twice with phosphate-buffered saline (PBS) at room temperature and then lysed in lysis T buffer (Peqlab Biotechnology GmbH, reference number 12-6634-01). Total RNA was then extracted with the peqGOLD Total RNA Kit C-Line (Peqlab Biotechnology GmbH, reference number: 12-6634-01) according to the manufacturer's instructions. RNA concentrations were determined by a Nanodrop spectrophotometer (ND-100 spectrophotometer; Nanodrop Technologies, Wilmington, DE, USA), and the RNA purity was ensured by a 260 / 280 ratio (mean 2.17 ± 0.04). For mRNA analysis, we synthesised cDNA using 2 µg of total RNA and qScript XLT cDNA SuperMix (Quantabio, product number 030256) as specified by the manufacturer and stored at -20°C. cDNA was amplified with a PerfeCTa qPCR fluorescent SYBR Green SuperMix (Quantabio, product number: Cat#95054-500) using real-time quantitative PCR (Rotor-Gene RG 6000— QIAGEN). The primer sequences used for the gene expression are shown in Table S1. To avoid amplification of contaminated genomic DNA, one of the two primers was placed at the junction between two exons. All relative gene expressions were quantified using the comparative Ct ($^{\Delta\Delta}$Ct) method [2] with Rplpo as the reference gene [3].

***Table S1*** *Primer sequences used for real-time polymerase chain reaction*

| Gene | Forward | Reverse |
|---|---|---|
| G6pdx | TGCACTTTGTCCGGAGTGAT | TGTGAGGGTTCACCCACTTG |
| Hk2 | CAAGTGCAGAAGGTTGACCA | GTGTGTGGTAGCTCCTAGCC |
| Oga | TGTGCAGTGGTTAGGGTGTC | GTCAGTGGGGGTGGTTGAAA |
| Ogt. | AGTGAAGGTGATGGCGGAAG | CCAGCCACATGGCCTGAATA |

| | | |
|---|---|---|
| Phgdh | TGGTGAACGCTAAGCTACTGG | CAGGGCCACAGTCAGGAG |
| Prps2 | GGCTGCGGGGAGATTAATGA | AATTGGAGCACGACTCTTGT |
| Rplpo | GGACCCGAGAAGACCTCCTT | GGACCCGAGAAGACCTCCTT |

G6pdx – Hexose-phosphate dehydrogenase; Hk2 – Hexokinase 2; Oga – *O*-GlcNAcase; Ogt – *O*-GlcNAc transferase; Phgdh – Phosphoglycerate dehydrogenase; Prps2 – Phosphoribosyl pyrophosphate synthetase 2; Rplp0 – Acidic ribosomal phosphoprotein P0 (reference gene); Shmt2 – Serine hydroxymethyltransferase-2.

**Protein determination**

After washing with ice cold PBS, cells were lysed on ice in 100 µl of radioimmunoprecipitation assay (RIPA) including 0.1% SDS,0.5 M sodium orthovanadate, 0.5% sodium deoxycholate, 50 mM NaF, 1 mM EDTA,150 mM NaCl, 1%Triton-X 100 and with protease and phosphatase inhibitor cocktail (Peqlab Biotechnology GmbH, Germany). Lysates were left on ice for 15 min and cellular debris removed by centrifuging at 13,000 rpm for 15 min, at 4°C. Supernatants were then transferred into fresh Eppendorf tubes and either frozen at -80°C for further Western blot analysis or to measure protein concentrations using the Bradford protein assay kit (BioRad Laboratories GmbH, Cat #5,000,111, Munich, Germany).

For the Phgdh experiment. protein synthesis in C2C12 myotubes was measured using SUnSET protein synthesis measurement as earlier [4]. Briefly, puromycin was added into cell culture medium to a final concentration of 1 µM exactly 30 min before harvesting the cells. Puromycin incorporation into proteins was measured by Western blot with anti-puromycin antibody (1:500, DSHB, Cat# PMY-2A4, RRID:AB_2619605). Total protein content was determined by using the bicinchoninic acid protein assay (Pierce, Thermo Scientific) with an automated Indiko plus analyzer (Thermo Scientific).

**Western blotting**

Protein samples containing 40 µg protein were solubilised in Laemmli sample buffer containing 10% β-mercaptoethanol and heated for 5 minutes at 95°C to denature proteins. Samples were then separated by electrophoresis in a 10% SDS-PAGE gel (Bio-rad, Germany). Proteins were transferred to polyvinylidene difluoride (PVDF) membranes (Biorad, Germany) using a Trans-Blot Turbo Blotting System (Bio-Rad, Germany). After this, a blocking step (5% non-fat milk powder, 1X Tris-buffered saline, 1% Tween-20) was performed, and the PVDF membrane was then incubated overnight at 4 °C with primary antibody against phospho-p70 S6 Kinase at Thr389 (1:1000, Cell Signaling Technology, #9205, RRID:AB_330944), mouse monoclonal anti-*O*-GlcNAc (1:5000, RL2; ThermoFisher Scientific; #MA1-072, RRID:AB_326364), and mouse anti-puromycin antibody (1:500, DSHB, Cat# PMY-2A4, RRID:AB_2619605). After incubating with horseradish peroxidase (HRP)-conjugated secondary anti-mouse (1:2000, Cell Signaling; #7076, RRID:AB_330924) or anti-rabbit IgG antibodies (1:2000, Cell Signaling, #7074, RRID:AB_2099233) for 1 h at room temperature, the membranes were developed with enhanced chemiluminescence (ECL) (Bio-Rad, Germany), and the signals were detected by an INTAS Chemocam Imager (Royal Biotech GmbH, Germany). Immunoreactive bands were quantified by using the ImageJ software (http://rsb.info.nih.gov/ij/index.html) and the strongest band of a Ponceau S stain of the membrane was used for normalisation [5]. In the case of the analysis of puromycin-incorporated proteins, total muscle *O*-GlcNAc-modified proteins, and total protein loading (stain-free), the intensity of the whole lane was quantified, and Western blot results of puromycin-incorporated proteins were normalised to the total protein loading analysed from the stain-free image of the membrane.

**Metabolomics analysis**

Muscle cells and plantaris muscle tissue were prepared based on [6]. The plantaris muscle tissue was inserted into an Eppendorf tube including 80% (vol/vol) high-performance liquid chromatography (HPLC)-grade methanol (cooled to −80 °C) in a ratio of 10 mg tissue to 0.2 mL extraction solvent, and then homogonised using a Precellys lysing kit (Bertin Instruments, USA) and a homogenizer machine (Bertin Technologies, USA) with high speed for 6 x 20 s and 1 minute rest in between, in which the samples were cooled on dry ice. Supernatant was transferred in new Eppendorf tubes, vortexed 2 x 20 s (1 minute rest in between, in which the samples were cooled on dry ice) and then incubated for 4 h at -80°C. After centrifugation at 14,000 g for 10 min at 4°C, the supernatant was transferred to a cryotube and stored at -80°C until further analysis.

Skeletal muscle cells were quickly washed with PBS. The PBS was carefully removed before the 6-well dish was placed on dry ice and quenched with 80% (vol/vol) HPLC-grade methanol (cooled to -80°C). After the cells were incubated for 20 min at -80°C, the dish was placed back on dry ice and the cells were scraped. The cell lysate/methanol mixture was transferred into a 1.5 ml tube and vortexed for 2x30 s at 4°C with 1 minute rest in between, in which the samples were cooled on dry ice. After incubation for 30 min at -80°C and centrifugation at 14,000 g for 5 min at 4°C, the metabolic-containing supernatant was transferred to cryotube and stored at -80°C until further analysis.

Untargeted metabolomics were performed using a Nexera UHPLC system (Shimadzu, Nakagyo-ki, Kyoto, Japan) coupled with a Q-TOF mass spectrometer (TripleTOF 6600, AB Sciex, Framingham, MA, USA). Separation of the samples was performed using a HILIC UPLC BEH Amide 2.1 × 100, 1.7 µm analytic column (Waters Corp., Milford, MA, USA), and a reversed-phase Kinetex XB-C18, 2.1 × 100, 1.7-µm analytical column (Phenomenex, Torrance, CA, USA), respectively. The mobile phase was 5 mM ammonium acetate in water (eluent A) and 5 mM ammonium acetate in acetonitrile/water (95/5, v/v) (eluent B) for HILIC separation and 0.1% formic acid (eluent A) and 0.1% formic acid (eluent B) in acetonitrile for reversed phase separation. The HILIC gradient profile was as follow: 100%B from 0 to 1.5 min, 60% B at 8 min and 20% B at 10 min to 11.5 min and 100% B at 12 to 15 min. The reversed-phase gradient profile was 0.2% B from 0 to 0.5 min to 100% B at 10 min which was held for 3.25 min. Afterwards the column was equilibrated to starting conditions. A volume of 5 µL per sample was injected for both HILIC and RP chromatographic methods. The autosampler was cooled to 10°C and the column oven heated to 40°C. A quality control (QC) sample which was pooled from all samples was injected after every 10th sample. Samples were measured in Information Dependent Acquisition (IDA) mode. MS settings in the positive mode were as follows: Gas 1 55, Gas 2 65, Curtain gas 35, Temperature 500°C, Ion Spray Voltage 5500 V, declustering potential 80 V. The mass range of the TOF MS and MS/MS scans were 50 - 2000 *m/z* and the collision energy was ramped from 15 - 55 V. MS settings in the negative mode were as follows: Gas 1 55, Gas 2 65, Cur 35, Temperature 500°C, Ion Spray Voltage -4500, declustering potential -80. The mass range of the TOF MS and MS/MS scans were 50 - 2000 *m/z* and the collision energy was ramped from -15 - -55 V.

The raw files were first converted to mzXML format via the ProteoWizard msConvert [7]. Peak-picking, alignment, correspondence performed using the peak-density method and peak-grouping were performed using the previously published R-package “xcms” available via the Bioconductor website for bioinformatic processing [8-10]. The ion mass of the features was compared to different databases (MS1 annotation), including the MS-DIAL DB, HMDB, PubChem and in-house reference DBs, allowing common modifications and a tolerance window of mass (15 ppm). The cosine distance between measured and reference spectra was calculated (MS2 annotation). The metabolite intensities were log transformed before statistical analysis. Detected metabolites were evaluated for correct annotation by reviewing chromatographic characteristics between DB and measured metabolites or by availability of data of previously measured reference standards. Analytical data of skeletal muscle cells were normalised to protein concentration using the Bradford method. The associated untargeted metabolomics data are available on the Mass Spectrometry Interactive Virtual Environment (MassIVE) data repository with ID MSV000090865.

Annotation of the metabolites detected by mass spectrometry were performed based on exact mass and by comparing the MS/MS-fragmentation pattern to databases. Unfortunately, we could not differentiate based on retention time, MS1 and fragmentation between glucose-6-phosphate, fructose-6-phosphate or glucose-1-phosphate which is why we took them together and referred to them as phosphorylated sugar. We could not distinguish between glucose and other hexoses and we referred to glucose. The same applies for ribose-5-phosphate and ribulose-5-phosphate, which was termed ribose-5-phosphate/ribulose-5-phosphate. N-Acetyl-D-Glucosamine-6-phosphate could not be distinguished from N-acetyl-D-mannosamine-6-phosphate and we referred to N-Acetylglucosamine-6-phosphate. We could not distinguish between UDP-N-acetylglucosamine (UDP-GlcNAc) and e.g. UDP-N-acetyl-D-mannosamine. Here, we referred to as UDP-N-acetylglucosamine as it is the second most abundant nucleotide-based structure in the cell after ATP.

#### [U-$^{13}C_6$]Glucose tracer experiment

To measure carbon incorporation in Igf-1 stimulated C2C12 myotubes, 4.5 g/L glucose consisting of ~20% [U-$^{13}C_6$]glucose and of 80% non-labelled glucose was added into glucose-free DMEM (Gibco, Cat#11966-025, Waltham, MA, USA) supplemented with 2 % horse serum (Sigma-Aldrich, H1138, NY, USA), and cells were incubated with this enriched media on day 3 post differentiation. After serum starvation (DMEM only) for 4 h, IGf-1 treatment (100 ng/ml) was added at the same time as the tracer, and cells were incubated for 24 h, after which the media was collected, and cells were harvested in 1 M perchloric acid (PCA) for stable isotope analysis (see below). Harvested cells were centrifuged (13,000 g, 10 min at 4°C) and the clear supernatant was transferred into new Eppendorf tubes and neutralised using potassium hydroxide. The pellet was washed twice with 70% (v/v) ethanol.

#### GC-MS analysis of [U-$^{13}C_6$]glucose

The pellet was hydrolysed overnight at 110°C in 1 ml of HCl (0.1 mol/ L) with 1 ml $H^+$ dowex resin in ddW. The hydrolysed free amino acids were isolated by passing through dowex columns, eluted into 2 mol/L $NH_4OH$ and dried down. The $^{13}C$ labelling of the protein-bound alanine (product) was determined using gas chromatography-combustion-isotope ratio mass spectrometry (GC-C-IRMS) following derivatization as its MCME derivative. The media alanine (precursor) enrichment is determined by gas chromatography mass spectrometry; briefly following deproteinization of 200ul media with 1ml ice cold ethanol, the supernatant is dried under nitrogen, reconstituted in 300ul 0.5M HCl and liquid-liquid extraction performed with 2ml Ethyl acetate to remove any lipid in the media. The aqueous fraction is dried then the amino acids derivatised as their t-BDMS derivative with MTBSTFA and the isotope ratio determined using single ion monitoring (SIM) of m/z 320 and 323.

The $^{13}C$ isotope ratios were converted to atom percent excess (APE), and the fractional synthesis rate (FSR) was calculated using the following equation: FSR (%/h) = $(APE_{product}/APE_{precursor})/t) * 100$ where $APE_{product}$ is the increase in protein-bound enrichment of $^{13}C$-alanine, $APE_{precursor}$ is the average in enrichment of $^{13}C$-alanine in the media and t = time in hours.

#### Bioinformatics analysis of human data

Publicly available summary statistics [11] from the published genome-wide association study (GWAS) on appendicular lean mass of UK Biobank participants (n=450,243) [12] were used to identify possible association ($p < 5.0e-8$) between single nucleotide polymorphisms (SNPs) located in or near genes involved in the regulation of the serine synthesis pathway and its downstream enzymes and appendicular lean mass. The UK Biobank is an open-access large prospective study with phenotypic and genotypic data from more than 500,000 participants (>90% of participants are of white ethnicity) with an age range for inclusion of 40–69 years when recruited in 2006–2010 [13].

In the next step, the Genotype-Tissue Expression (GTEx) portal [14] was used to analyze the association between the lean mass related SNPs and expression of specific genes in different tissues ($p < 0.05$). The GTEx project is an ongoing effort to build a comprehensive public resource to study tissue-specific gene expression and regulation. Samples were collected from 49 tissue sites across >800 individuals, primarily for molecular assays including whole genome sequencing (WGS), whole exome sequencing (WES), and RNA-Seq. SNPs that were significantly ($p < 0.05$) correlated with expression of genes were considered as expression quantitative trait loci (eQTLs).

#### Statistical Analysis

One-way ANOVA followed by Tukey HSD was applied for the post-hoc analysis using Prism version 8.0 statistical software package (GraphPad Prism; RRID: SCR_002798) for the comparison of multiple groups with statistical significance set at $p<0.05$. The unpaired two-tailed Student's t-test was performed for analysing protein synthesis after Phgdh treatment and for metabolomics analysis to compare the average of two groups. The reported unadjusted p values for all metabolites in the cell culture study were subsequently adjusted to account for multiple hypotheses testing. The false discovery rate (FDR) method of Benjamini and Hochberg was used to perform the adjustment (FDR<0.1). We did not find significant results in the *in vivo* study after adjusting for multiple hypotheses testing FDR<0.1. We therefore took metabolites into account of the *in vivo* study, which were significant after adjusting for multiple hypotheses testing FDR<0.2 *in vivo*, and were also significant in the *in vitro* experiment. Results were expressed as mean ± standard error (SEM).

#### Systematic Review

To place the results of the in *vivo* and *in vitro* experiments in context, we performed a systematic review regarding metabolic changes during muscle hypertrophy and added the experimental results to the systematic review. We registered this systematic review on PROSPERO (access number: CRD42022318998; (https://www.crd.york.ac.uk/PROSPERO/). To identify metabolites that changes their concentration during muscle growth, we carried out a systematic review according to the PRISMA guidelines. One researcher searched in PubMed, Scopus and metabolite using the PICO framework [15] leading to the following search strategy: ((mice or mouse) and (metabolomics or metabolome or metabolomes) and (skeletal muscle)) and ((increase or increased or increases or increasing or increasingly) or (mass gain) or (hypertrophy)) until May 11, 2022. Two researchers then screened the search results. Discrepancies were resolved by mutual decisions regarding the individual studies. Please find the retracted information of the individual publications in Table 1. We included articles from peer-reviewed journals, written in English, that performed metabolomics analysis in both both mouse *in vivo* and skeletal muscle cell *in vitro* studies [using resistance/strength training-related exercises, skeletal muscle overload intervention, electrical pulse stimulation, stretch-induced skeletal muscle hypertrophy, hypertrophy-related genetical manipulation techniques (including knockdown, skeletal muscle overload, knockout, knock-in and overexpression techniques) and pharmaceutical interventions]. We excluded studies where one or more of the following applied:

for animal studies:

- Other animals than mice
- Humans
- Pathological models (diabetes, cancer)
- Xenografts
- Death after gene manipulation
- *ex vivo* studies
- *In silico* studies

for *in vitro* studies:

- Non-skeletal muscle cell lines
- Pathological genetically modified skeletal muscle cells (e.g., diabetes, cancer)

for both animal and *in vitro* studies:

- No (exercise) intervention to induce skeletal muscle hypertrophy, or no skeletal muscle hypertrophy-related gene modification
- Only nutritional intervention to induce muscle hypertrophy
- Immune therapy
- No outcome measure of skeletal muscle hypertrophy

Two researchers individually assessed the quality of each individual study using the 10-item checklist of CAMARADES (Collaborative Approach to Meta Analysis and Review of Animal Experimental Studies) (Table S2). A metabolite set enrichment analysis was performed using the MetaboAnalyst 5.0 web platform [16]. We, alongside other researchers [17], faced challenges distinguishing between glucose-6-phosphate, fructose-6-phosphate, or glucose-1-phosphate, collectively termed phosphorylated sugar. In the systematic review, however, phosphorylated sugar was defined as glucose-6-phosphate for a better illustration. Importantly, glucose-6-phosphate was detected and significant in two other studies [18, 19], so that this assembly did not alter the overall outcome of integrating glucose-6-phosphate into the systematic review.

***Table S2*** *The Collaborative Approach to Meta-Analysis and Review of Animal Data from Experimental Studies (CAMARADES) checklist for study quality.*

| **Author (year)** | **year** | **(1)** | ***(2)*** | **(3)** | **(4)** | **(5)** | **(6)** | **(7)** | **(8)** | **(9)** | **(10)** | **Total score** |
|---|---|---|---|---|---|---|---|---|---|---|---|---|
| Baumert et al. | 2022 | Y | Y | NA | NA | N | Y | Y | N | Y | Y | 6 |
| Cheng et al. | 2014 | Y | Nk | N | NA | N | Y | Y | N | Y | Y | 5 |
| Collins-Hooper et al. | 2015 | Y | Y | N | NA | N | Y | Y | N | Y | Y | 6 |
| Jiang et al. | 2021 | Y | Nk | Y | Nk | N | Y | Y | Nk | Y | Y | 6 |
| Weyrauch et al. | 2020 | Y | Y | Y | NA | N | Y | Y | N | Y | Y | 7 |
| Wu et al. | 2017 | Y | Nk | N | NA | N | Y | Y | N | Y | Y | 5 |

Studies fulfilling the criteria of: (1) peer reviewed publication; (2) control of temperature; (3) random allocation to treatment or control; (4) allocation concealment; (5) blinded assessment of outcome; (6) avoidance of anesthetics with marked intrinsic properties; (7) animal model (aged, diabetic, or hypertensive); (8) sample size calculation; (9) compliance with animal welfare regulations; and (10) statement of potential conflict of interests. Also see supplementary material. Nk, not known

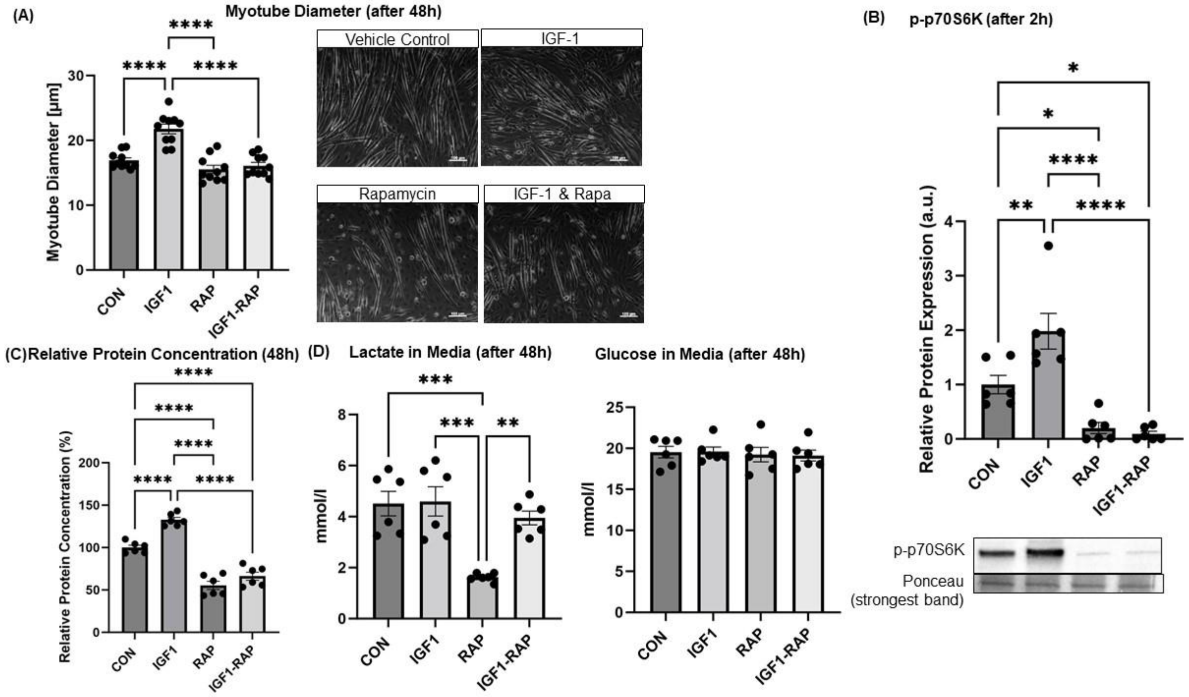


***Figure S1*** *(A) Myotube diameter measured 48 h after initialisation of treatments, (B) the phosphorylation of 70-kDa ribosomal protein S6 kinase (p70S6K) measured 2 h after initialisation of treatments. (C) Relative protein concentration normalised to vehicle control 48 h after initialisation of treatments. (D) Lactate and glucose concentration in conditioned media after 48 h. *Significant differences between indicated conditions, one-way ANOVA with Tukey HSD post-hoc test (p<0.05). Data are expressed as mean ± standard error (SEM).*

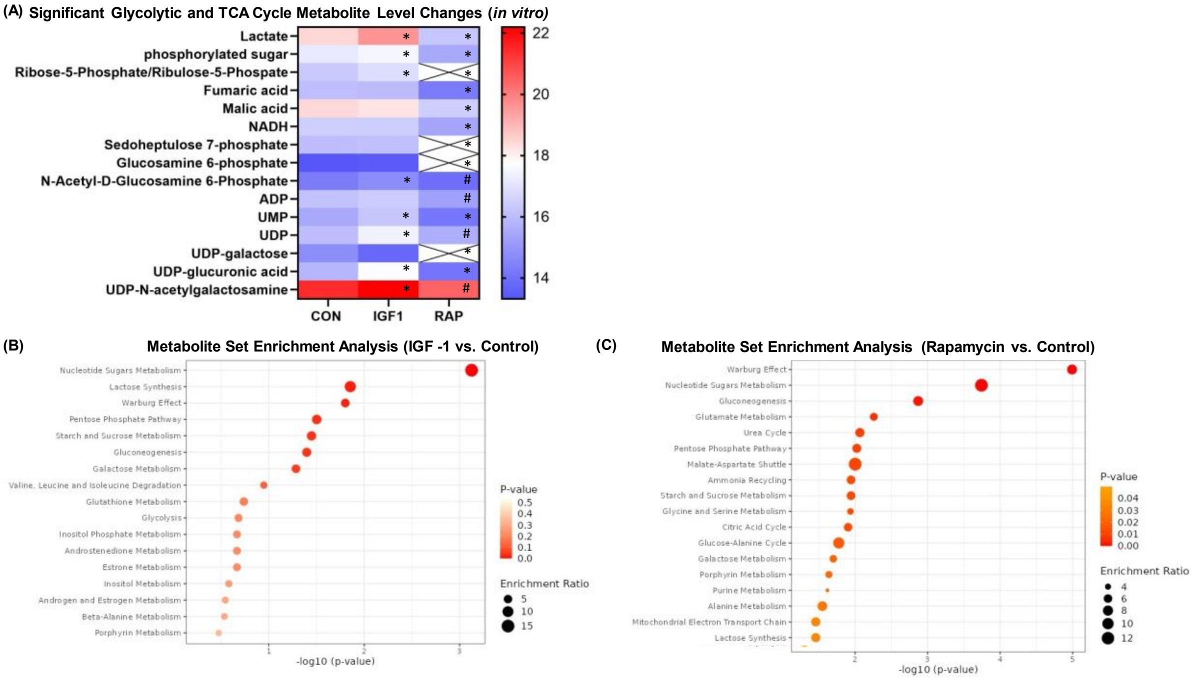


**Figure S2** *(A) Heatmap of significant clycolytic metabolite level changes after vehicle control, IGF-1 or rapamycin treatment in vitro. Crosses indicate that the metabolite is below the detection limit. Crosses indicate that the metabolite is below the detection limit. *Significant differences between treatment (IGF-1 or Rapamycin) compared to vehicle control; # Significant differences between IGF-1 compared to Rapamycin. (B) Enrichment analysis with MetaboAnalyst 5.0 between IGF-1 and control treatment and (C) rapamycin and control treatment. ATP = Adenosine diphosphate; CON = Control; IGF1 = Insulin-like growth factor-1; NADH = Nicotinamide adenine dinucleotide (reduced); RAP = Rapamycin; TCA = Tricarboxylic Acid Cycle; UMP = Uridine monophosphate; UDP = Uridine diphosphate.*

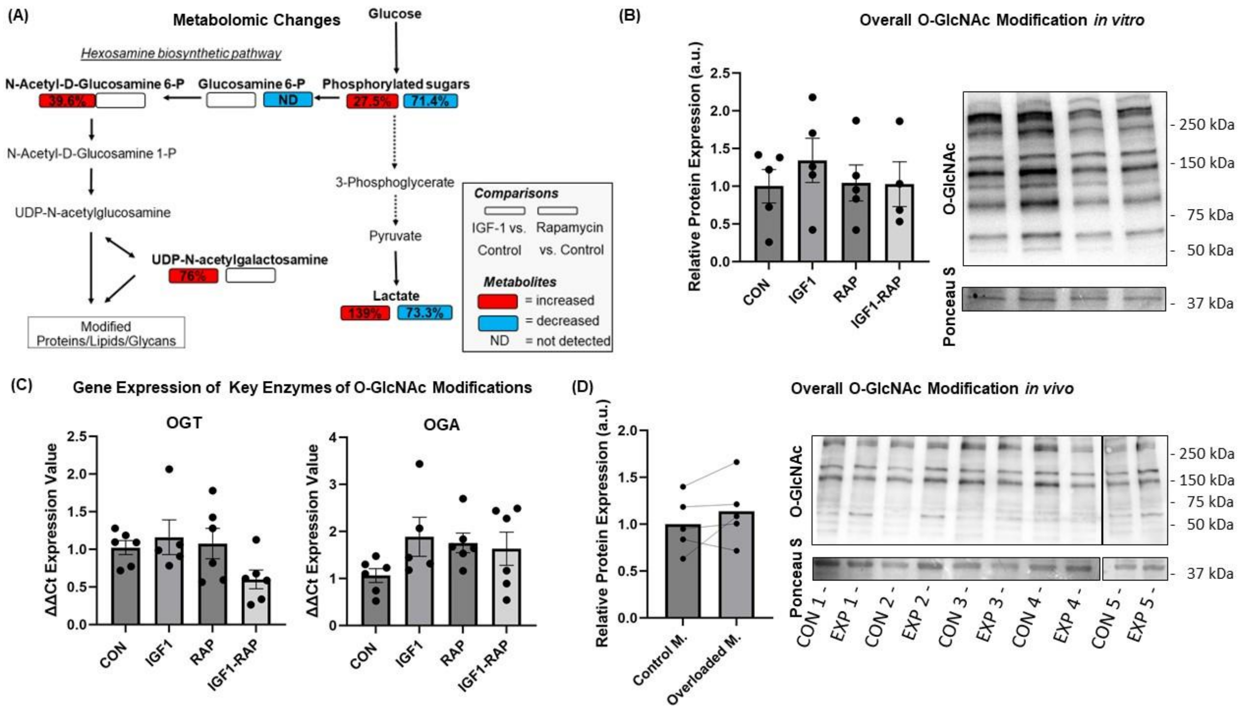


***Figure S3** Metabolomic changes in differentiated myotubes treated with either vehicle control, Insulin-like growth factor-1 (IGF-1; 100 ng/mL) or rapamycin (10 ng/mL) for 48 h (n = 3). Metabolites highlighted in bold were detected by untargeted metabolomics. Left box illustrates IGF-1 vs. control and right box rapamycin vs. control. Boxes highlighted in red and blue show significantly increased and decreased metabolite intensity levels, respectively (both FDR<0.2). White box shows no significant changes (FDR≥0.2). (B) Relative protein expression of total muscle O-N-acetylglucosamine (O-GlcNAc)-modified proteins normalised to vehicle control of the in vitro experiment after 48 h. O-GlcNAc antibody was normalised to the strongest band of a Ponceau S stain. (C) Gene expression of key enzymes of the post-translational modification termed O-linked N-acetylglucosaminylation of protein O-GlcNAc transferase (Ogt) and protein O-GlcNAcase (Oga) after 48 h. (D) Relative Protein expression of total muscle O-GlcNAc-modified proteins normalised to vehicle control of the in vitro experiment after 48 h. O-GlcNAc antibody was normalised to the strongest band of a Ponceau S stain. CON = Control; IGF-1 = Insulin-like growth factor-1; ND = Not detected; P = Phosphate; RAP = Rapamycin; TCA = Tricarboxylic acid cycle; UDP = Uridine diphosphate. *Significant differences between groups, one-way ANOVA with Tukey HSD post-hoc test (p<0.05). Data are expressed as mean ± standard error (SEM).*

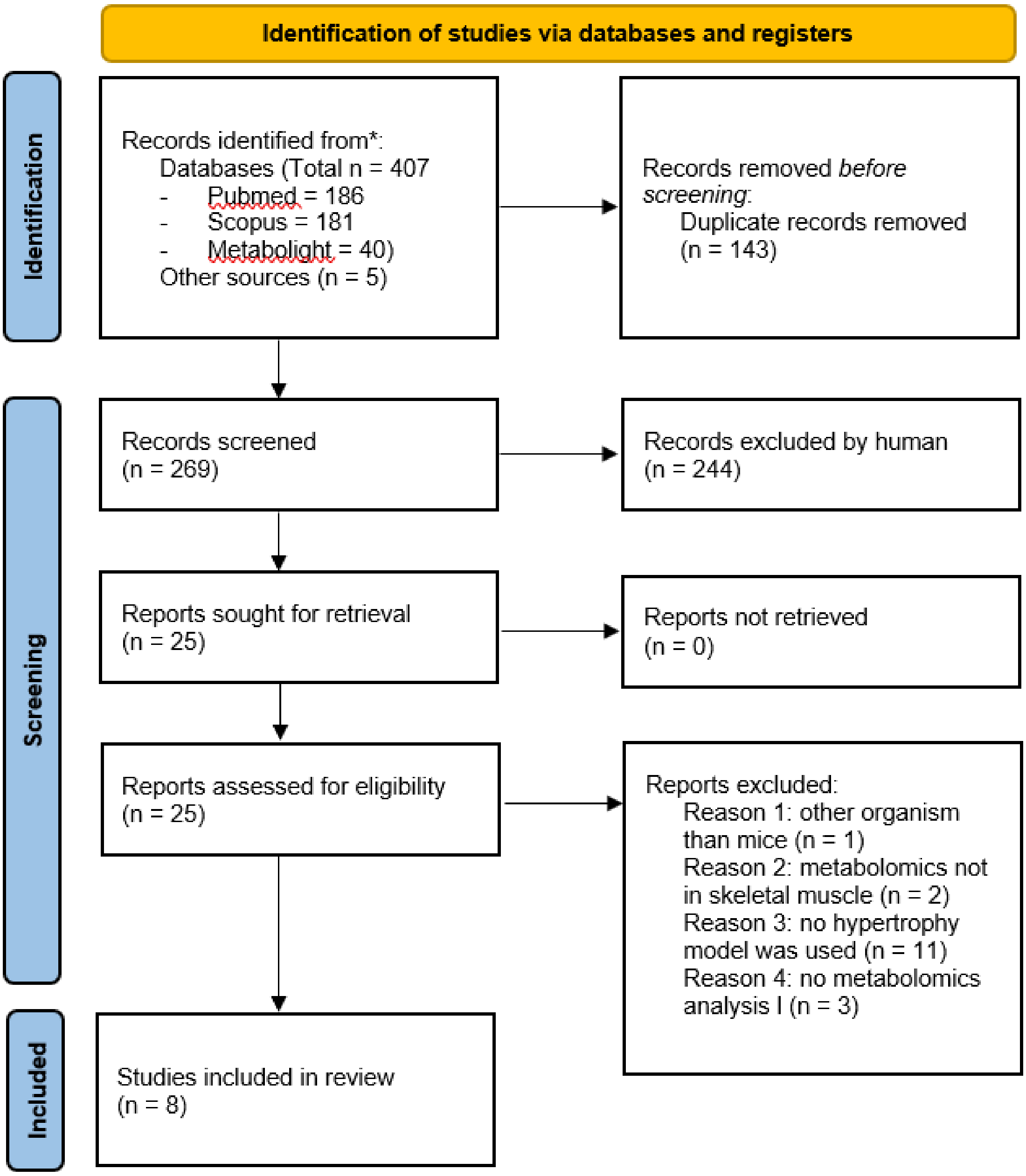


***Figure S4*** *PRISMA 2020 flow diagram for new systematic reviews which included searches of databases and registers only*
